# Multi-tier signaling and epigenomic reprogramming orchestrate microglial inflammatory states and functions associated with demyelination

**DOI:** 10.1101/2025.08.21.670551

**Authors:** Félix Distéfano-Gagné, Nesrine Belhamiti, William Saxon, Sara Bitarafan, André Machado Xavier, Stéphanie Fiola, Serge Rivest, David Gosselin

## Abstract

The extensive heterogeneity of microglia inflammatory states accompanying neurodegenerative diseases underscores the complex molecular mechanisms that regulate these cells. Here, we report on transcriptional mechanisms that control microglial inflammatory polarization associated with brain demyelination in mice. Using flow cytometry, microscopy and RNA-seq, we identified two dominant, functionally distinct states inflammatory microglia: Clec7a+CD229+CD11c-microglia that are prone to proliferation and that transcribe high levels of extracellular matrix-associated genes, and Clec7a+CD229+CD11c+ microglia characterized by inflammatory tissue-remodeling and antigen presentation gene signatures. Epigenomic analyses implicated genome-wide nucleosome remodeling to the polarization process, driven by state-associated hierarchal stimulation of pro-inflammatory transcription factors, as well as re-calibration of Mef2 homeostatic input. Mechanistically, we confirmed relevance for Trem2, Mef2a and Egr2 to the microglial inflammatory polarization and demyelinating processes. Therefore, distinct configurations of signaling input cooperate with epigenetic mechanisms to reprogram the transcriptional output of microglia to enable their inflammatory functions.

## Introduction

The ability to deploy and coordinate inflammatory responses in the central nervous system (CNS) is a defining feature of microglia, the resident macrophages of the CNS parenchyma^1-3^. Through stimulation of immune and stress-response signaling pathways, inflammatory microglia will transcribe the gene effectors required for proliferation, enhanced endocytic and cargo processing capabilities, and antigen presentation, to name a few functions^4^. However, the overall microglial population at lesioned sites is comprised of several distinct subsets of microglia that each display subset-biased inflammatory gene and protein signatures, which confer each subset with an overall distinct ensemble of cellular capabilities^5-14^. While it remains unclear whether these distinct gene signatures also endow a subset with unique, non-redundant functions, such heterogeneity points to substantial complexity in the regulatory activity of the signaling and transcriptional effectors in coordinating gene expression in microglia.

Evidence supports a role for a growing number of ligands and receptors in promoting microglial inflammatory subset polarization and activity in chronic brain lesions. For instance, activation of the Trem2 receptors drives the “disease-associated microglia” (DAM) polarization process^5,15,16^. Stimulation of the IFNAR receptor complex contributes to the type 1 interferon transcriptional response displayed by microglia in demyelinating injuries^17^. Additional receptors implicated include Il-3r, Il-6ra, Tnfr1 and purinergic receptors, to name a few^18-21^. Upon ligand binding, these receptors will stimulate myriads of intracellular signaling cascades, culminating with the activation of signal-dependent transcription factors (SDTFs) that will deliver regulatory input to the transcriptional machinery at the genomic level^22^. To date, a few SDTFs have been proposed to regulate microglial transcriptional response in degenerative brain lesions. For example, Stat1 and Stat3 activity promotes pro-inflammatory, neural-damaging, activity of microglia, while Irf5, Lxr and Mitf activity increases transcription of effector genes involved in debris clearance, phagolysosomal functions and cholesterol processing^23-28^. Alternatively, Bhlhe40/41 may provide negative feedback on the induction of genes involved in lipids processing^23^. But owing to the wide array of receptors feeding inflammatory input to microglia in lesions, as well as the extensive collection of families of pro-inflammatory SDTFs expressed in these cells, our understanding of inflammatory gene regulation in these cells remains rudimentary.

Here, we sought to identify and characterize the transcriptional regulators that specify the inflammatory cell states and functions of mouse microglia associated with chronic demyelination, a pathology linked with MS. To this end, we used the cuprizone (CPZ) model of demyelination, in which dietary administration to mouse of cuprizone neurotoxin triggers oligodendrocyte cell death and demyelination throughout the brain^29^. Notably, a robust inflammatory response driven by microglia accompanies lesion development, and the quality of this response impacts neural damage burden and repair^30,31^. Through analyses of transcriptional signatures and genome-wide chromatin states, coupled with loss-of-function experiments, our analyses revealed that pan-acting transcriptional effectors cooperate with subset-biased input to specify polarization states and functions of distinct inflammatory microglia subsets. Together, these analyses shed light on multiple intricate layers of transcriptional regulation necessary to orchestrate complex inflammatory activity of microglia in chronic brain lesions.

## Results

### Protein expression patterns of Clec7a, CD229 and CD11c reveal functionally distinct microglial states in the demyelinating brain

We first characterized the composition of the microglial population associated with brain demyelinating lesions. For this, we used flow cytometry to phenotype CD11b^+^CD45^+^CD44^Low/Neg^Ace^Low/Neg^ microglia isolated from mice fed CPZ diet for 4 weeks (CPZ-4w), interrogating cell surface expression of Clec7a, CD229, and CD11c proteins to discriminate among possible subset (Figure S1A); these were selected based on their reported association with the DAM and MGnD transcriptional signatures^5,32^. Overall, each of those markers displayed elevated expression in CPZ-4w microglia compared to healthy brain microglia. Magnitude of increase was not uniform however, with upregulated expression of Clec7a, CD11c and CD229 respectively labeling on average 25%, 18% and 50% of the global microglia population (Figure 1A). Though the spectrum of gain of Clec7a expression was relatively restrained, microscopy analyses confirmed detection of Clec7a+ cells in the brain of CPZ-4w mice, but not in that of healthy mice (Figure S1B). Co-expression analyses revealed that CD229+ microglia exhibited higher expression of Clec7a compared to their CD229-counterparts and that Clec7a expression was homogenous within the CD229+ subset (Figure 1B). In contrast, expression levels of CD11c partitioned CD229+ microglia into CD11c- and CD11c+ states (Figure 1C). Confocal microscopy confirmed that CD11c expression may discriminate distinct subsets of Iba+Clec7a+ phagocytes, as both Iba1+Clec7a+CD11c+ and Iba1+Clec7a+CD11c-microglia were readily detected in the brain of mice fed CPZ for 4 weeks (Figures 1D and S1C). No CD11c+ signal was detected in the healthy brain, coherent with the flow cytometry data (Figure S1B). While we were not able to identify an antibody to reveal CD229+ microglia in the demyelinating brain, these data show that CD11c expression discriminated at least 2 major subsets of inflammatory microglia in the demyelinating brain.

**Figure 1.**
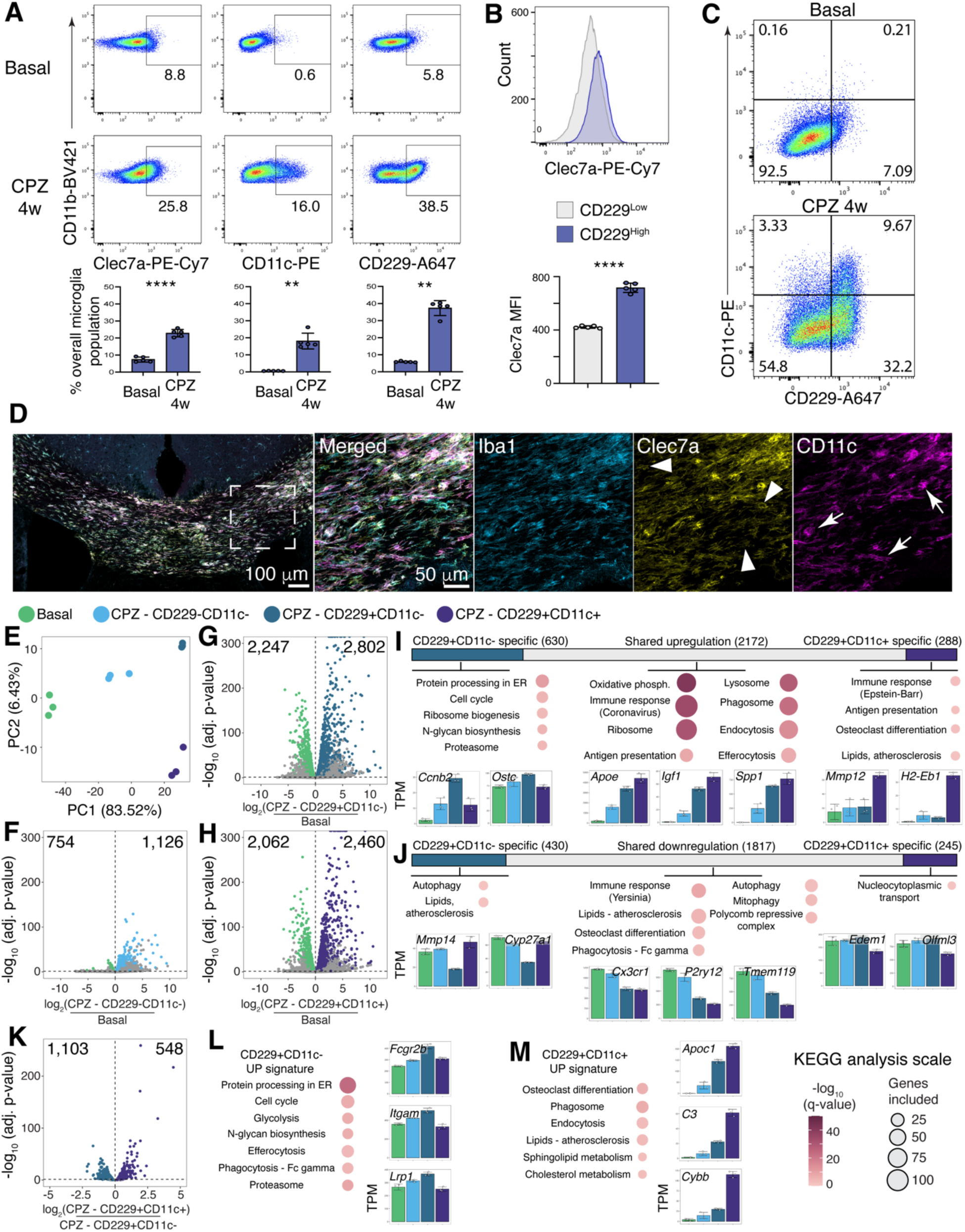
Identification of distinct microglial inflammatory states in the demyelinating brain. **(A)** Flow cytometric assessment of Clec7a, CD229 and CD11c expression on healthy and CPZ-4w microglia (Clec7a normal distribution, unpaired Welch’s t-test, two-tailed; CD11c and CD229 not normally distributed, unpaired Mann-Whitney t-test, two-tailed, *n* = 5). **(B)** Flow cytometric quantification of Clec7a median fluorescence intensity for CPZ-4w CD229- and CD229+ microglia (normal distribution, unpaired Welch’s t-test, two-tailed; n = 5). **(C)** Flow cytometry identification of CD229-CD11c-, CD229+CD11c-, CD229+CD11c+, and CD229-CD11c+ microglia subsets in mice fed CPZ for 4 weeks. **(D)** Representative confocal microphotograph depicting expression of Clec7a and CD11c on Iba1+ cells in the brain of mice fed CPZ for 4 weeks. Arrowheads depict Iba1+Clec7a+CD11c-cells, while arrows depict Iba1+Clec7a+CD11c+ cells. **(E)** Principal component analysis of RNA-seq data from homeostatic and CPZ-4 microglia subsets (*n* = 3). **(F, G, H)** Volcano plots of RNA-seq data comparing gene profiles from homeostatic with CPZ-4w microglia subsets. Differentially expressed genes are colored coded (DESeq2, FDR ≤ 0.05; *n* = 3). **(I, J)** Bar chart depicting shared and subset-specific upregulation (I) or downregulation (J) in CD229+CD11c- and CD229+CD11c+ CPZ-4w microglia (see G, H). Dominant KEGG terms associated and bar graphs depicting expression of representative genes are annotated (Transcripts Per Million (TPM) normalization). **(K)** Volcano plots of RNA-seq data depicting expression of genes from (I), comparing expression in CPZ CD229+CD11c+ to CPZ CD229+CD11c-microglia. Differentially expressed genes are colored coded (DESeq2, FDR ≤ 0.05; *n* = 3). **(L, M)** Dominant KEGG terms associated with genes preferentially expressed in CPZ CD229+CD11c- vs CPZ CD229+CD11c+ (L), or vice versa (M), and bar graphs depicting mRNA expression of representative genes (TPM, linked to K). All bar graphs are means ± S.D.; ∗p < 0.05, ∗∗p < 0.01, ∗∗∗p < 0.001, ∗∗∗∗p < 0.0001.

Next, we performed RNA-seq analyses to infer the potential cellular competences of the main inflammatory subsets of microglia in the CPZ brain, as defined by CD229 and CD11c expression. Using fluorescence-activated cell sorting (FACS), we isolated CD229-CD11c-, CD229+CD11c- and CD229+CD11c+ microglia subsets from CPZ-4w mice and homeostatic microglia from healthy mice (e.g., CD11b+CD45^+^CD44^Low/Neg^ Ace^Low/Neg^ cells). Our strategy efficiently excluded non-microglial cell types, as indicated by expression of markers for border-associated macrophages, monocytes, astrocytes, oligodendrocytes, neurons, endothelial and ependymal cells that fell below threshold of expression, e.g., 16 TPM (Table S1). Principal component (PC) analysis indicated that both CD229+ subsets were the most distinct compared to homeostatic microglia; molecular events linked to the CPZ diet and gain of CD229 expression were responsible for 83.25% of the variance observed (Figure 1E). Overall, 3,090 and 2,492 genes were respectively significantly upregulated and downregulated in at least one of the two CD229+ subsets compared to microglia from the healthy brain microglia (false discovery rate (FDR) of 0.05; transcripts per million (TPM) ≥ 16; Figure 1F to 1J and Table S1). Genes induced in both CD229+ microglia subsets (2,172) included *Apoe*, *Igf1* and *Spp1*; based on KEGG pathway analysis, several of those genes contribute to oxidative phosphorylation, immune defense and otherwise general macrophage functions such as endocytosis and phagolysosomal activity (Figure 1I). In contrast, 630 genes were exclusively upregulated in CD229+CD11c-cells, and these comprised effectors of cell proliferation (e.g., *Ccnb2*), protein processing, and biosynthesis of ribosome and N-glycans (e.g., *Ostc*) (Figure 1I); flow cytometry validated that the CD229+CD11c-subset comprises the largest proportion of proliferative Ki67+ microglia (Figure S1D). Lastly, 288 genes displayed increased expression specifically in the CD11c+ subset; these comprised additional mediators of immune defense and antigen-presentation (*Mmp12*, *H2-Eb1*) (Figure 1I).

We next sought to define more precisely each subset’s gained specialized functions relative to one another. For this, we recovered the genes among the global set of 3,090 upregulated genes defined above that each CD229+ subset express at higher levels compared to its counterpart, at a FDR of 0.05 (Figure 1I). Overall, 1,103 and 548 genes were more highly expressed in the CD11c- and CD11c+ subsets, respectively (Figure 1K and Table S1). While the gene signatures recovered were aligned with those of the preceding analyses, results also suggested that each subset may engage in internalization and processing of molecular cargos through different mechanisms. For example, CD229+CD11c-microglia may be more apt at engaging in efferocytosis and phagocytosis through Fc gamma, owing to their higher expression of genes such as *Fcgr2b*, *Itgam* and *Lrp1* (Figure 1L). In contrast, the CD11c+ microglia express at higher levels a battery of genes involved in endocytosis and cholesterol metabolism, including *Apoc1* and *C3*, along with superoxide-generating enzyme *Cybb* (Nox2) (Figure 1M). Finally, individual inspection of differentially expressed genes suggested that these subsets contribute non-redundantly to the inflammatory and tissue-remodeling processes associated with active demyelination. For example, CD229+CD11c-expressed highest levels of proteoglycan *Cspg4*, extracellular matrix glycoprotein *Fn1*, regulator of metalloproteinase *Bsg* (EMPRINN), serine protease *Htra3*, and cytokines *Vegfa* and *Vegfb* (Figure S1E). Alternatively, CD229+CD11c+, exhibited stronger expression of metalloproteinases *Mmp12* and *Mmp14* (Figures 1I and 1J), secreted extracellular matrix protein *Postn*, and upregulation of chemokine *Ccl5* and cytokines *Il1b and Pdgfa* (Figure S1E).

Overall, these data agree with previous studies that reported heterogeneity of microglial states and functions in lesioned brains. They also extend, however, on current literature by suggesting that high protein expression of CD229 and Clec7a may broadly define lesion-associated microglia, and that CD11c co-expression defines a more select subset of inflammatory microglia.

### Candidate regulators instructing microglial inflammatory subsets in the demyelinating brain

Next, we investigated the signaling and transcriptional effectors that coordinate the polarization of the different microglia subsets in the demyelinating brain. For this, we analyzed the genome-wide profiles of chromatin accessibility associated with each state, as revealed by ATAC-seq^33^. As immune and several other SDTFs interact with the genome predominately at promoter-distal cis-regulatory genomic elements (dCREs)^22,34^, we focused our analyses on accessible areas that fall outside the -1000 to 500 bp window of annotated transcriptional start sites. Coherent with RNA-seq data, PC analysis of ATAC-seq data from FACS-sorted microglia showed that both CPZ CD229+ subsets are most distinct from the healthy brain microglia, and that CPZ CD229+ -linked variables drive most of the variance associated with these data (Figure 2A). Overall, several thousand promoter-distal regions underwent local chromatin remodeling across the CPZ subsets (Figure 2B). Most of these were contributed by either CD229+ subsets, and paired comparisons of these two subsets showed that they shared relatively similar genome-wide profiles of chromatin accessibility. Indeed, a total of 1,244 areas were differentially accessible, with a sub-group of 643 sites displaying increased accessibility, and another 601 being less accessible, in CD229+CD11c+ microglia compared to CD229+CD11c-microglia (Figures 2B and 2C).

**Figure 2.**
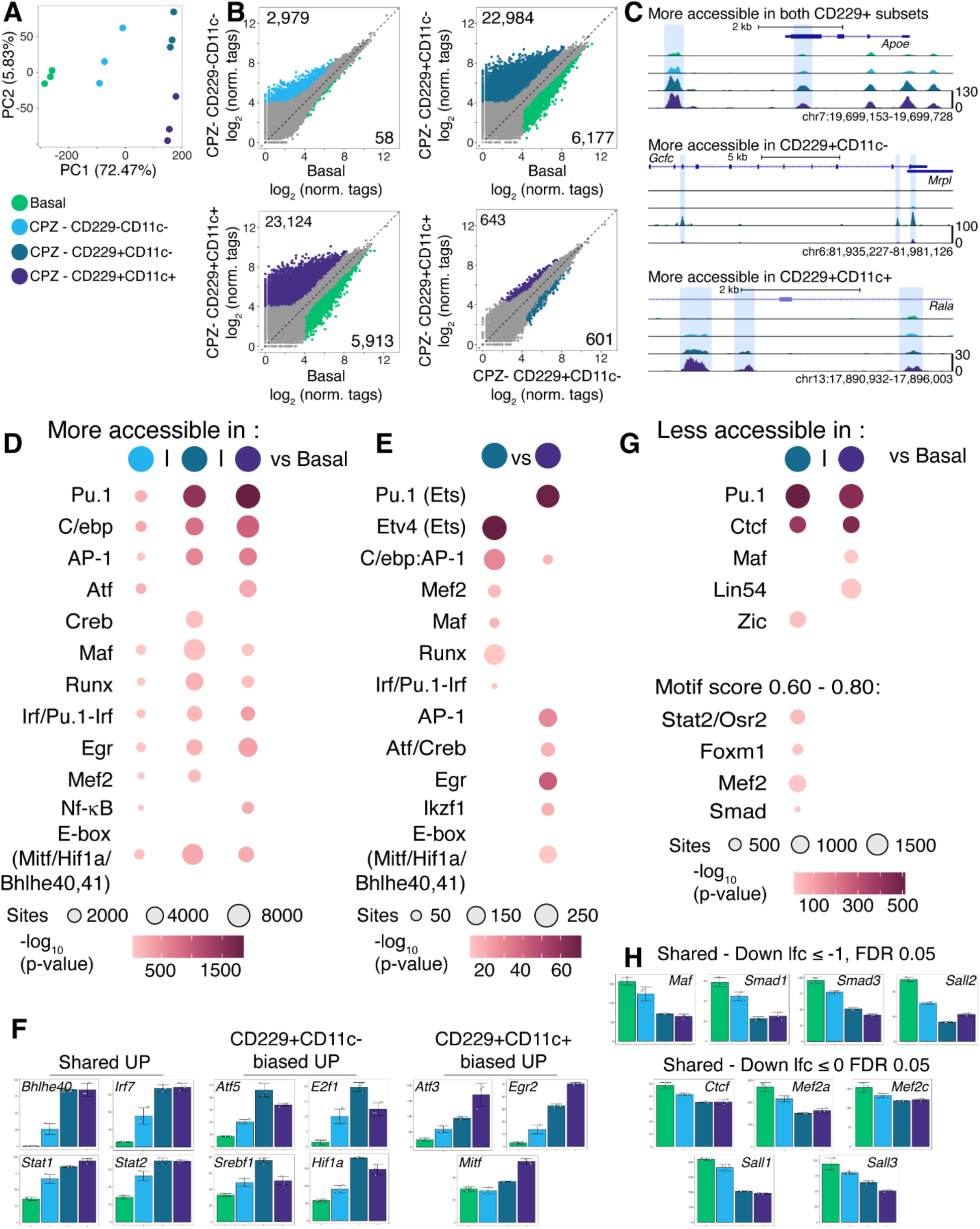
Inflammatory microglial states in the demyelinating brain display distinct profiles of chromatin accessibility. **(A)** Principal component analysis of ATAC-seq data from homeostatic and CPZ-4 microglia subsets (*n* = 3). **(B)** Scatterplots of ATAC-seq data, centered on promoter-distal genomic sites, comparing profiles from homeostatic and CPZ-4w microglia subsets. Sites differentially accessible are colored coded (DESeq2, FDR ≤ 0.05; *n* = 3). **(C)** UCSC genomic browser displays of ATAC-seq data of representative loci that exhibit significantly higher accessibility in CPZ CD229+CD11c-, CD229+CD11c+, or both subsets, compared to homeostatic microglia. **(D, E)** De novo DNA motifs enrichment analysis for promoter-distal genomic sites that gain accessibility in CPZ CD229+ microglia vs homeostatic microglia (D), or in CPZ CD229+CD11c- vs CPZ CD229+CD11c+ and vice versa (E). **(F)** mRNA levels (TPM) of select transcription factors significantly upregulated in either or both CPZ CD229+ subsets compared to homeostatic microglia (DESeq2, FDR ≤ 0.05; log_2_ fold change ≤ 1). **(G)** De novo DNA motifs enrichment analysis for promoter-distal ATAC-seq sites that lose chromatin accessibility in CPZ CD229+CD11c- or CPZ CD229+CD11c+ vs homeostatic microglia. **(H)** mRNA levels (TPM) of select transcription factors significantly downregulated in either or both CPZ CD229+ microglia subsets compared to homeostatic microglia. (DESeq2, FDR ≤ 0.05; log_2_ fold change ≤ 0 or -1). All bar graphs are means ± S.D.

De novo motifs enrichment analyses performed on sites that gain accessibility in each CPZ subsets compared to microglia from the healthy brain identified similar families of TFs as potential regulators of microglial immune activity in the demyelinating brain. These included Pu.1, AP-1, C/ebpb, Atf, Pu.1-Irf/Irf, Runx, Maf, Mef2, Nf-κB, and E-box binding factors, which may include Hif1, Mitf, and Bhlhe family members (Figure 2D). However, the number of accessible sites with enrichment for these motifs varied across the subsets, suggesting that characteristics pertaining to the quality (i.e., different family members, dimers) and/or strength of input may vary in a subset-dependent manner. This was also supported by direct comparison of the CD229+ subsets with one another. Here, promoter-distal sites more accessible in CD229+CD11c-displayed stronger enrichment for Runx, Maf and Mef2, while those more accessible in CD229+CD11c+ may be subject to stronger input from Atf and Egr family members, as well as E-box binding factors, among others.

We then interrogated the RNA-seq datasets to identify TFs whose regulatory function could in part be controlled at the level of mRNA expression. At a FDR of 0.05, a total of 36 and 75 were upregulated and downregulated, respectively, in at least one of the CD229+ subsets (Table S2). Fold changes in gene expression, however, varied greatly among these, ranging from log_2_ fold-change (lfc) of 0.09 to 7.4 for upregulated TFs. Applying a two-fold threshold to changes in expression levels retained 15 and 15 *robustly* upregulated and downregulated factors, again respectively (Figures S2A, S2B and Table S2). Both CD229+ subsets upregulated 4 TFs to similar extent: *Bhlhe40*, *Irf7, Stat1* and *Stat2* (Figures 2F and S2A). Of interest, the CD229+CD11c-subset expressed 8 TFs at significantly higher levels compared to CD11c+ microglia. These included *Atf5*, *Hif1a* along with TFs previously implicated with cell proliferation in microglia such as *E2f1*, *Etv4* and *Srebf1*, which is coherent with the enhanced proliferative gene signature of this subset^35^. Alternatively, CD229+CD11c+ microglia preferentially upregulated *Atf3*, *Egr2* and *Mitf*.

ATAC-seq data analysis also revealed robust enrichment for Pu.1 and Ctcf motifs at sites that lose accessibility in CPZ CD229+ subsets, but otherwise very modest enrichment for additional factors for either subset (Figure 2G). That said, motifs partially overlapping with Mef2 and Smad were significantly enriched at sites less accessible in the CD229+CD11c-subset, which would be coherent with context-dependent re-calibration and/or down-tuned activated for these factors. RNA-seq data analyses also corroborated this: CD229+ subsets downregulated mRNA expression of *Mef2a*, *Mef2c*, several *Smad*s, and *Ctcf*, as well as that of members of the Sall family of transcriptional co-repressor (Figures 2H and S2B).

Together, these data suggest that overlapping but ultimately distinct hierarchies of pro-activating inflammatory input interface with homeostatic signals to promote the subset-biased inflammatory states and functions of microglia in the demyelinating brain.

### Graded dependence of CD229+ inflammatory subsets polarization on Trem2 signaling

We next performed loss of function experiments to gain insight into the signaling mechanisms that control the TFs implicated in microglia inflammatory subset polarization identified above. To this end, previous studies showed that several microglial inflammatory gene signatures associated with chronic brain lesions arise through Trem2 receptor-dependent and -independent signaling^5,36^. Thus, to assess how the polarization of CD229+ microglia subsets depends on Trem2 signaling, we examined microglial inflammatory states in WT and Trem2^-/-^ mice after 4 weeks of CPZ feeding. Flow cytometric analysis showed that absence of Trem2 completely impeded emergence of CD229+CD11c+ microglia and severely compromised induction of CD229+CD11c- and proliferative Ki67+ microglia (Figures 3A to D). Microscopy analyses corroborated these observations. For example, while Iba1 signal surface coverage over white matter regions, irrespective of intensity, was similar in CPZ WT and Trem2^-/-^, Trem2-null mice displayed a 50% reduction and a near complete absence in Clec7a+ and CD11c+ staining surface coverage, respectively (Figure 3E). Lastly, the altered microglial response in Trem2^-/-^ mitigated demyelination in the white matter (Figure 3F). Together, these results confirm that the polarization process of distinct microglia inflammatory phenotypes rely to different degrees on Trem2 signaling.

**Figure 3.**
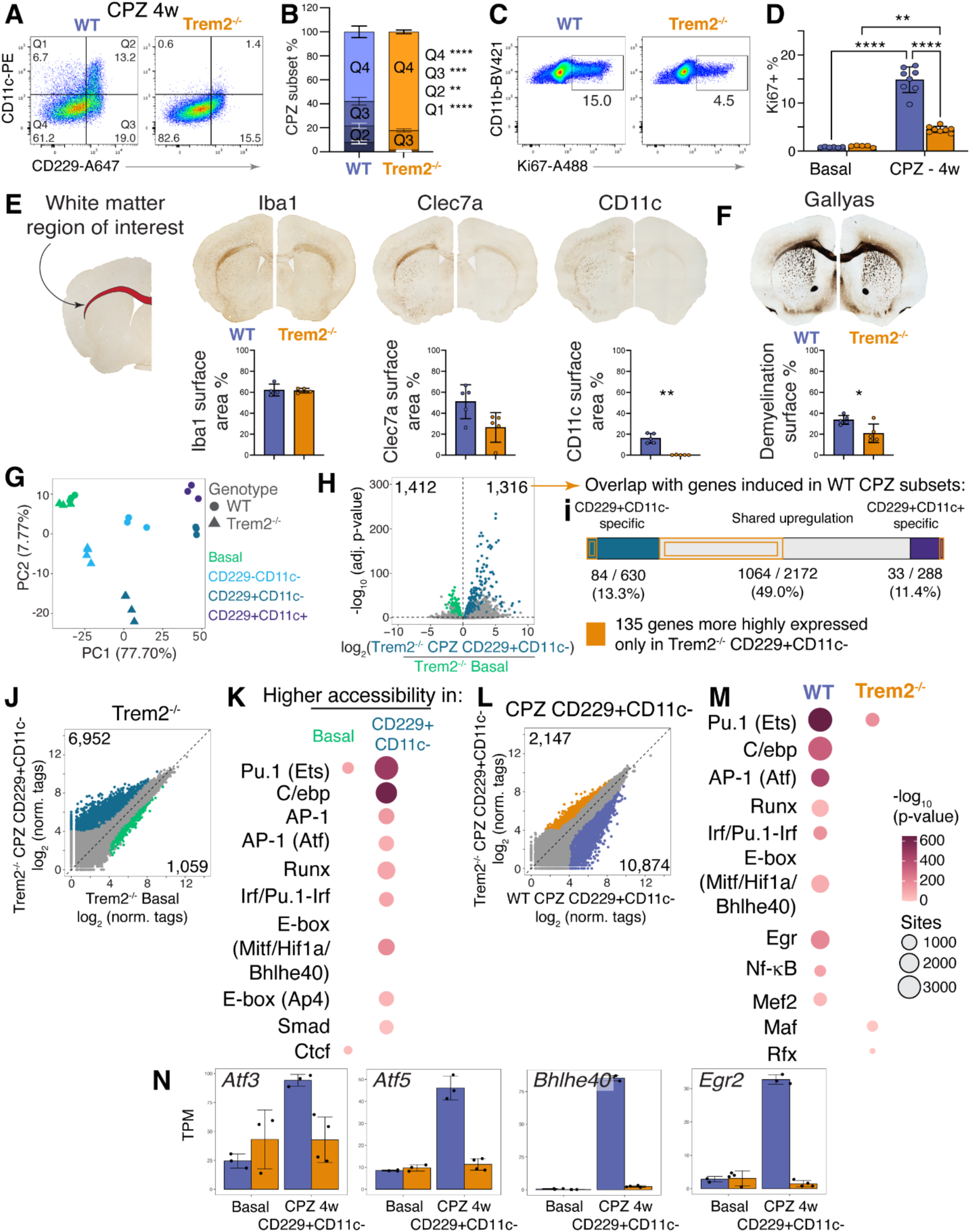
Trem2-independent and -dependent regulatory input promote polarization of CD229+ inflammatory microglial subsets. **(A, B)** Flow cytometric assessment (A) and quantification (B) of CPZ microglia subset from WT and Trem2^-/-^ mice (Multivariate ANOVA (MANOVA) followed by a Games-Howell post-hoc test; *n* = 7-8). **(C, D)** Flow cytometry scatterplots (C) and quantification (D) of Ki67+ proliferative microglia in the brain of WT and Trem2^-/-^ mice fed regular or CPZ diet for 4 weeks (two-way ANOVA followed be Tukey’s multiple comparison test) **(E, F)** Representative brain microphotographs depicting Iba1, Clec7a and CD11c immuno-labeling (E), and modified Gallyas staining (F), comparing WT and Trem2^-/-^ mice fed CPZ for 4 weeks. Appended below, quantification of positive signal (E) or negative (F) signal coverage over corpus callosum and external capsule (Iba1, CD11c, Gallyas: normal distribution, unpaired Welch’s t-test, two-tailed; Clec7a: not normal distribution, Mann-Whitney unpaired t-test two-tailed; One sample value (WT - Iba1 met criterion for “outlier” and was removed (see Method details); *n* = 4-5). **(G)** Principal component analysis of RNA-seq data from homeostatic and CPZ microglia subsets, from WT and Trem2^-/-^ mice (*n* = 4). **(H)** Volcano plot of RNA-seq data comparing homeostatic and CPZ CD229+CD11c-microglia from Trem2^-/-^ mice. Differentially expressed genes are colored coded (DESeq2, FDR ≤ 0.05; *n* = 3-4). **(I)** Bar chart displaying number of genes upregulated in Trem2^-/-^ CPZ CD229+CD11c-, defined in (H), that overlap with subset-shared and -specific upregulated gene signature in WT CPZ CD229+ subsets, defined in figure 1I. **(J)** Scatterplots of ATAC-seq data, centered on promoter-distal genomic sites, comparing profiles from homeostatic and CPZ CD229+CD11c-microglia from Trem2^-/-^ mice. Sites differentially accessible are colored coded (DESeq2, FDR ≤ 0.05; *n* = 2-3). **(K)** De novo DNA motifs enrichment for promoter-distal genomic sites that gain or lose chromatin accessibility in Trem2^-/-^ CPZ CD229+CD11c- vs Trem2^-/-^ homeostatic microglia. **(L)** Scatterplots of ATAC-seq data, centered on promoter-distal genomic sites, comparing profiles from Trem2^-/-^ CPZ CD229+CD11c-microglia to WT CPZ CD229+CD11c-. Sites differentially accessible are colored coded (DESeq2, FDR ≤ 0.05; *n* = 2-3). **(M)** De novo DNA motifs enrichment analysis for promoter-distal genomic sites that gain or lose chromatin accessibility in Trem2^-/-^ CPZ CD229+CD11c-microglia vs WT counterpart (linked to L). **(N)** mRNA levels (TPM) of select transcription factors expressed at significantly lower levels in Trem2^-/-^ CPZ CD229+CD11c- vs WT CPZ CD229+CD11c-microglia. All bar graphs are means ± S.D. ∗p < 0.05, ∗∗p < 0.01, ∗∗∗p < 0.001, ∗∗∗∗p < 0.0001.

RNA-seq analyses extended impact of Trem2-dependent and independent signaling to the regulation of distinct gene programs. First, principal component analysis of RNA-seq data for subsets isolated from WT and Trem2^-/-^ revealed that under basal condition, microglia from both genotypes are highly similar (Figure 3G); direct comparison revealed that only 10 genes are differentially expressed (Figure S4A and Table S3). However, robust divergences occurred in the context of CPZ-induced demyelination, despite CD229+CD11c-microglia arising in both genotypes. For example, CPZ CD229+CD11c- from Trem2^-/-^ upregulated 1,316 genes compared to healthy state Trem2^-/-^ microglia (Figure 3H and Table S3). However, only 84 of these genes overlapped with the CPZ-4w CD229+CD11c- -specific genes signature from WT microglia, as defined in figure 1j (Figure 3I). Instead, a majority of these genes, i.e., 1,064/1,316, overlapped at 49% with the *shared* signature of induced genes (Figure 3I). Comparison of CD229+CD11c- microglia from both genotypes with one another further underscored the importance of Trem2 signaling to proper gene regulation in these cells in the demyelinating brain; 2,504 and 1,799 genes were, respectively, expressed at significantly lower and higher levels in Trem2^-/-^ CD229+CD11c- microglia relative to their WT counterparts (Figure S3A and Table S3). Therefore, Trem2-independent input may play key roles in inducing general inflammatory gene programs, but they act relatively weakly on subset-biased gene program regulation.

To gain insights into the transcriptional effectors of Trem2-dependent and -independent regulatory mechanisms, we performed ATAC-seq centered on CPZ CD229+CD11c- microglia from Trem2^-/-^ mice. First, these displayed gains and losses of chromatin accessibility at 6,952 and 1,059 promoter-distal sites, respectively, relative to homeostatic, basal Trem2^-/-^ microglia (Figure 3J). DNA motif enrichment analysis revealed that motifs linked to Ets, AP-1, C/ebp, Runx, Irf and E-box-binding factors are highly enriched within the 6,952 sites that gained accessibility in CD229+CD11c- (Figure 3K), thus connecting regulation of these factors to Trem2-independent signaling. Second, contrasting the Trem2^-/-^ CPZ CD229+CD11c- subset to its WT counterpart revealed that while 2,148 sites gained accessibility in Trem2^-/-^, 10,874 sites were more accessible in WT (Figure 3L). Notably, sites more accessible in the WT subset displayed enrichment for Egr, Nf-κB and Mef2 factors, in addition to Ets, AP-1, C/ebp, Runx, Irf and E-box-binding factors included in the preceding comparison (Figure 3M). Of interest, differential gene expression analyses indicated the lower activity signature for several of these factors may be linked to failure of Trem2^-/-^ microglia to induce their expression at the transcriptional level in the demyelinating brain. In particular, Trem2^-/-^ CPZ CD229+CD11c- failed to upregulate *Bhlhe40*, *Atf3*, *Atf5*, *Egr2*, *Etv4*, *E2f1* and *Maff* compared to homeostatic microglia (Figures 3N and S3C). These cells also conserved homeostatic-like expression level for *Ctcf*, and decreased to lesser extent expression of *Mef2a*, *Smad1, Smad3, Sall1* and *Sall3* compared to WT CPZ CD229+CD11c- microglia (Figure S3D).

Collectively, these data suggest that Trem2-mediated upregulation of transcription factors such as *Bhlhe40*, *Atf3* and *Egr2* may be critical for inflammatory microglia to fully achieve the CD229+CD11c- state and commit to the CD11c+ phenotype. They also suggest that pan-acting immune regulators, including Pu.1, AP-1, C/ebp, Runx and Irf family members, along with limited recalibration of homeostatic input such as Mef2 and Sall factors, can coalesce in a Trem2-independent manner to induce a partial CD229+CD11c- state that is associated, among other, with lesion-triggered proliferative activity of microglia.

### Decreased Mef2a activity in microglia initiates CD229+CD11c- state polarization in the healthy brain

We next tested the hypothesis that downregulation of a defined homeostatic transcriptional input may facilitate CD229+CD11c- state polarization. For this, we performed loss-of-functions experiments, targeting Mef2a on the basis of 1) DNA motifs analyses that suggested loss of Mef2 input at sites more accessible in WT compared to CD229+ (Figure 2G), 2) its high expression in homeostatic microglia that undergoes downregulation in CPZ microglia (Figure 2H), and 3) milder downregulation of *Mef2a* mRNA in Trem2^-/-^ CPZ CD229+ microglia compared to WT (Figure S4D). To inhibit Mef2a expression in adult microglia, we generated Cx3cr1^CreERT2/WT^:Mef2a^fl/fl^ mice, hereafter referred to as MG-Mef2aΔ. Cx3cr1^CreERT2/WT^:Mef2a^WT/WT^ were used as controls (i.e., Ctrls), and both genotypes underwent the same Tamoxifen injection protocol^37,38^. Overall, this strategy led to a ∼90% reduction in expression of *Mef2a* transcripts that include the targeted exon in microglia from MG-Mef2aΔ mice relative to controls (Figure 4A). Such decrease in *Mef2a* mRNA expression had a relatively mild impact on gene expression in microglia however, with only 49 and 35 genes being significantly less or more highly expressed, respectively, in whole MG-Mef2aΔ microglia compared to Ctrl (Figure 4B and Table S4). Furthermore, both Ctrl and MG-Mef2aΔ mice had similar Iba1+ microglial cell density in cortical regions (Figure 4C). That said, flow cytometry detected a significant ∼2.5-fold increase in the proportion of CD229+ microglia in the brain of MG-Mef2aΔ mice (Figure 4D). Proportion and expression level of Clec7a augmented slightly as well, but proportions of CD11c+ or Ki67+ microglia were not different (Figure 4D). RNA-seq analyses revealed that CD229+ microglia expressed at higher levels 20 genes compared to its CD229- complement; these comprised several genes expression in microglia in lesioned context, including *Apoe*, *Axl*, *Ctse*, and *Lyz2*, to name a few (Figures 5E, 5F and Table S4). Lastly, ATAC-seq revealed that 431 promoter-distal sites lose accessibility in microglia from MG-Mef2aΔ mice compared to microglia from Ctrls (Figure 4G). Of note, these sites displayed enrichment for motifs linked to Pu.1 and Mef2 factors (Figure 4H). The 108 sites that gained accessibility in MG-Mef2aΔ did not exhibit enrichment for any particular motifs. Both CD229+ and CD229- microglia in MG-Mef2aΔ mice exhibited similar patterns of chromatin accessibility at non-promoter genomic regions (Figure 4I).

**Figure 4.**
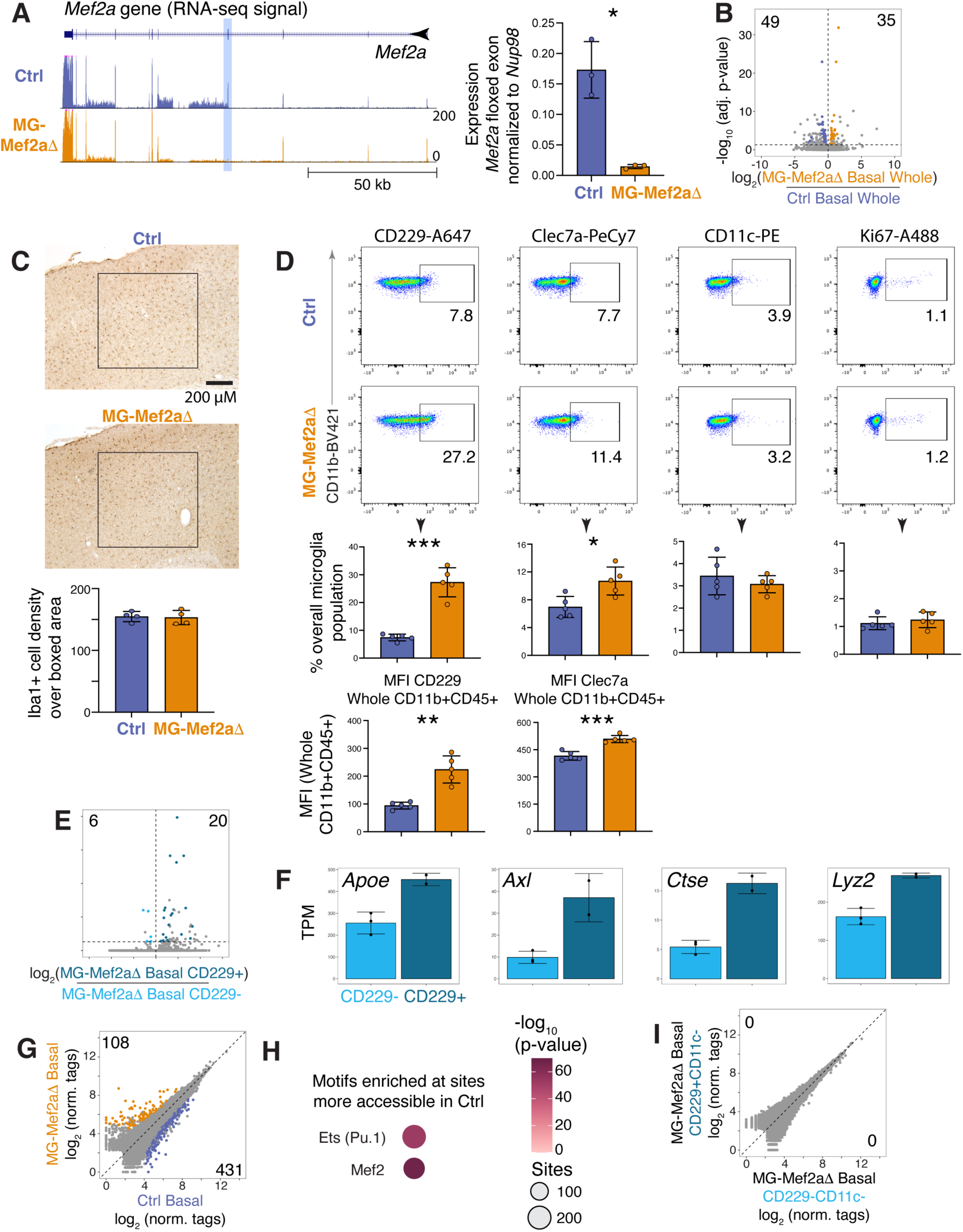
Decreased *Mef2a* mRNA expression in microglia promotes CD229+ polarization in the homeostatic brain. **(A)** Left: UCSC genome browser display of RNA-seq data at *Mef2a* locus, comparing homeostatic Ctrl vs MG-Mef2aΔ microglia 8 weeks after Tamoxifen injection. *Mef2a* floxed exon highlighted in blue. Right: *Mef2a* floxed exon RNA-seq reads count normalized to reads count over *Nup98* gene body (normally distributed, unpaired Welch’s t-test, two-tailed; *n* = 3). **(B)** Volcano plot of RNA-seq data, comparing WT vs MG-Mef2aΔ whole microglia 8 weeks after Tamoxifen injection. Differentially expressed genes are colored coded (DESeq2, FDR ≤ 0.05; *n* = 3). **(C)** Representative brain microphotographs and quantification of Iba1+ microglia, in cortical region of healthy male Ctrl and MG-Mef2aΔ mice (normal distribution, unpaired Welch’s t-test, two-tailed). **(D)** Flow cytometric assessment of CD229, Clec7a, CD11c and Ki67 expression for microglia from Ctrl and MG-Mef2aΔ mice. Below, quantifications of subset proportion and median fluorescence intensity (CD229, Clec7a, CD11c are normally distributed, unpaired Welch’s t-test, two-tailed; Ki67 not normally distributed, Mann-Whitney test; *n* = 5). **(E)** Volcano plot of RNA-seq data comparing CD229+CD11c- vs CD229-CD11c- microglia from healthy MG-Mef2aΔ mice. Differentially expressed genes are colored coded (DESeq2, FDR ≤ 0.05; *n* = 2-3). **(F)** mRNA levels (TPM) of representative genes significantly upregulated in CD229+CD11c- relative to CD229-CD11c- microglia (linked to E). **(G)** Scatterplots of ATAC-seq data, centered on promoter-distal genomic sites, comparing WT vs MG-Mef2aΔ whole microglia 8 weeks after Tamoxifen injection. Differentially accessible sites are colored coded (DESeq2, FDR ≤ 0.05; *n* = 2-4). **(H)** De novo DNA motifs enrichment analysis for promoter-distal sites with higher chromatin accessibility in Ctrl whole microglia compared to MG-Mef2aΔ. **(I)** Scatterplots of ATAC-seq data, centered on promoter-distal genomic sites, comparing CD229+CD11c- to CD229-CD11c- microglia isolated from healthy MG-Mef2aΔ mice. No sites are differentially accessible (*n* = 2). All bar graphs are means ± S.D. ∗p < 0.05, ∗∗p < 0.01, ∗∗∗p < 0.001.

**Figure 5.**
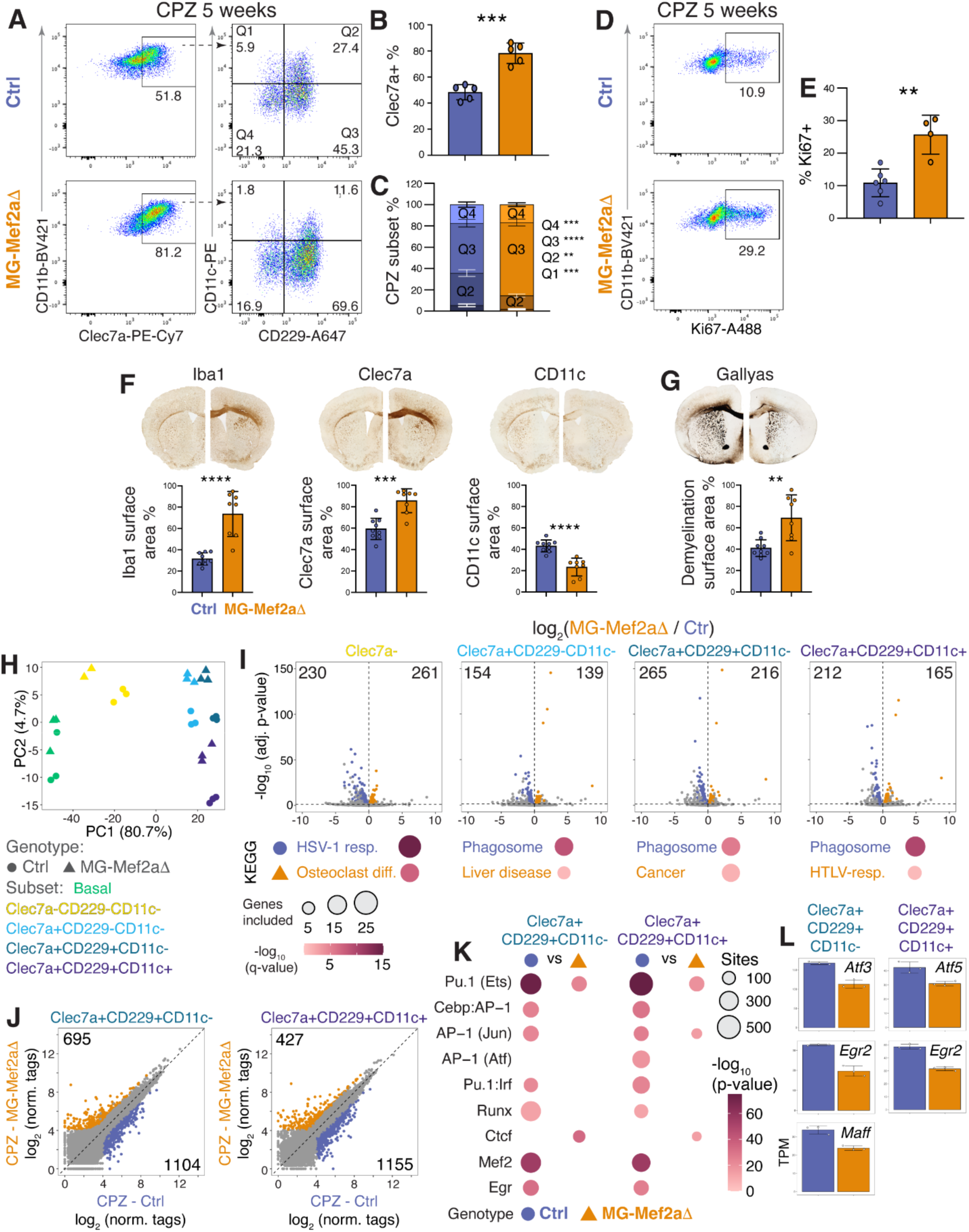
Decreased *Mef2a* mRNA expression in microglia exacerbates neuroinflammation and myelin damage in mice fed cuprizone diet. **(A)** Flow cytometric assessment of Clec7a, CD229 and CD11c expression for microglia from Ctrl and MG-Mef2aΔ mice fed CPZ for 5 weeks (*n* = 5). **(B)** Quantification of Clec7a+ microglia proportion (linked to A; normal distribution, unpaired Welch’s t-test, two-tailed; *n* = 5). **(C)** Quantification of proportion of CPZ microglia subsets, defined by CD229 and CD11c expression, linked to A (Q1 to Q4; multivariate ANOVA (MANOVA) followed by a Games-Howell post-hoc test; *n* = 5). **(D, E)** Flow cytometry scatterplots (C) and quantification (D) of proliferative microglia inflammatory subsets, revealed by Ki67, in the brain of Ctrl and MG-Mef2aΔ mice fed CPZ diet for 5 weeks (normal distribution unpaired Welch’s t-test, two-tailed; *n* = 4-6) **(F, G)** Representative brain microphotographs depicting Iba1, Clec7a, and CD11c immuno-labeling (F), and modified Gallyas staining (G), comparing Ctrl and MG-Mef2aΔ mice fed CPZ for 5 weeks. Below, quantification of positive (F) or negative (G) area coverage over corpus callosum and external capsule (Iba1, CD11c, Gallyas: normal distribution, unpaired Welch’s t-test, two-tailed; Clec7a: not normal distribution, Wilcoxon-Mann-Whitney, unpaired t-test- two-tailed; *n* = 8-9). **(H)** Principal component analysis of RNA-seq data from microglia subsets from Ctrl and MG-Mef2aΔ mice fed regular or CPZ for 5 weeks. **(I)** Volcano plots of RNA-seq data comparing CPZ microglia subsets from Ctrl and MG-Mef2aΔ mice. Differentially expressed genes are colored coded (DESeq2, FDR ≤ 0.05; *n* = 3). Below, dominant KEGG term for differentially expressed genes in each subset. **(J)** Scatterplots of ATAC-seq data, centered on promoter-distal genomic sites, comparing MG-Mef2Δ CPZ CD229+CD11c- and CD229+CD11c+ microglia subsets to their Ctrl counterparts. Sites differentially accessible are colored coded (DESeq2, FDR ≤ 0.05; n = 3-4). **(K)** De novo DNA motifs enrichment analyses performed on promoter-distal genomic sites that display higher or lower chromatin accessibility in MG-Mef2Δ CPZ CD229+CD11c- and CD229+CD11c+ microglia subsets compared to their Ctrl counterparts. **(L)** mRNA expression levels (TPM) of select transcription factors that are expressed at significantly lower levels in MG-Mef2aΔ CPZ CD229+CD11c- and CPZ CD229+CD11c+ microglia compared to their Ctrl microglia counterparts (linked to I) All bar graphs are means ± S.D.; ∗p < 0.05, ∗∗p < 0.01, ∗∗∗p < 0.001, ∗∗∗∗p < 0.0001.

Overall, these results indicate that constant Mef2a input in microglia is necessary to counteract an intrinsic propensity to deploy inflammatory gene programs. As such, these data support a mechanism whereby initiation of inflammatory functions may not always depend on detection of pro-inflammatory stimulus, if a down-tuning in homeostatic input occurs.

### Decreased Mef2a activity in microglia amplifies inflammation and damage in the demyelinating brain

The observations above suggested that proper Mef2a activity in homeostatic microglia may counter these cells’ hardwired predisposition to engage in inflammatory functions; a similar interpretation was previously made regarding Mef2c activity in microglia^39^. However, the actual functional relevance of these gatekeeping mechanisms to the development and/or progression of chronic brain pathologies has yet to be validated. To address this, we compared CPZ-induced brain inflammatory responses and myelin damage burden of MG-Mef2aΔ vs Ctrl mice. As shown in Figure 5A, inhibiting Mef2a activity in microglia disrupted the abundance and relative configuration of inflammatory microglia subsets in the demyelinating brain, after 5 weeks of CPZ diet. Specifically, Clec7a+ and CD229+CD11c- accumulated in greater proportion in the brain of MG-Mef2aΔ mice, while relative frequency of CD229+CD11c+ decreased (Figures 5A, 6B and 6C). Assessment of Ki67 expression suggested that absence of Mef2a activity also potentiated the proliferative activity of microglia (Figures 5D and 6E). Microscopy analyses corroborated that loss of Mef2a input exacerbates microglial inflammatory activity associated with demyelination, as supported by the significant increase in Iba1+ and Clec7a+ immunostaining in white matter regions of MG-Mef2aΔ mice (Figure 5F). Of note however, we observed a decrease in CD11c+ signal intensity, which also aligns with flow cytometry observations above (Figure 5F). Finally, these alterations were associated with greater demyelination in MG-Mef2aΔ mice, as measured by modified Gallyas staining (Figure 5G).

**Figures 6.**
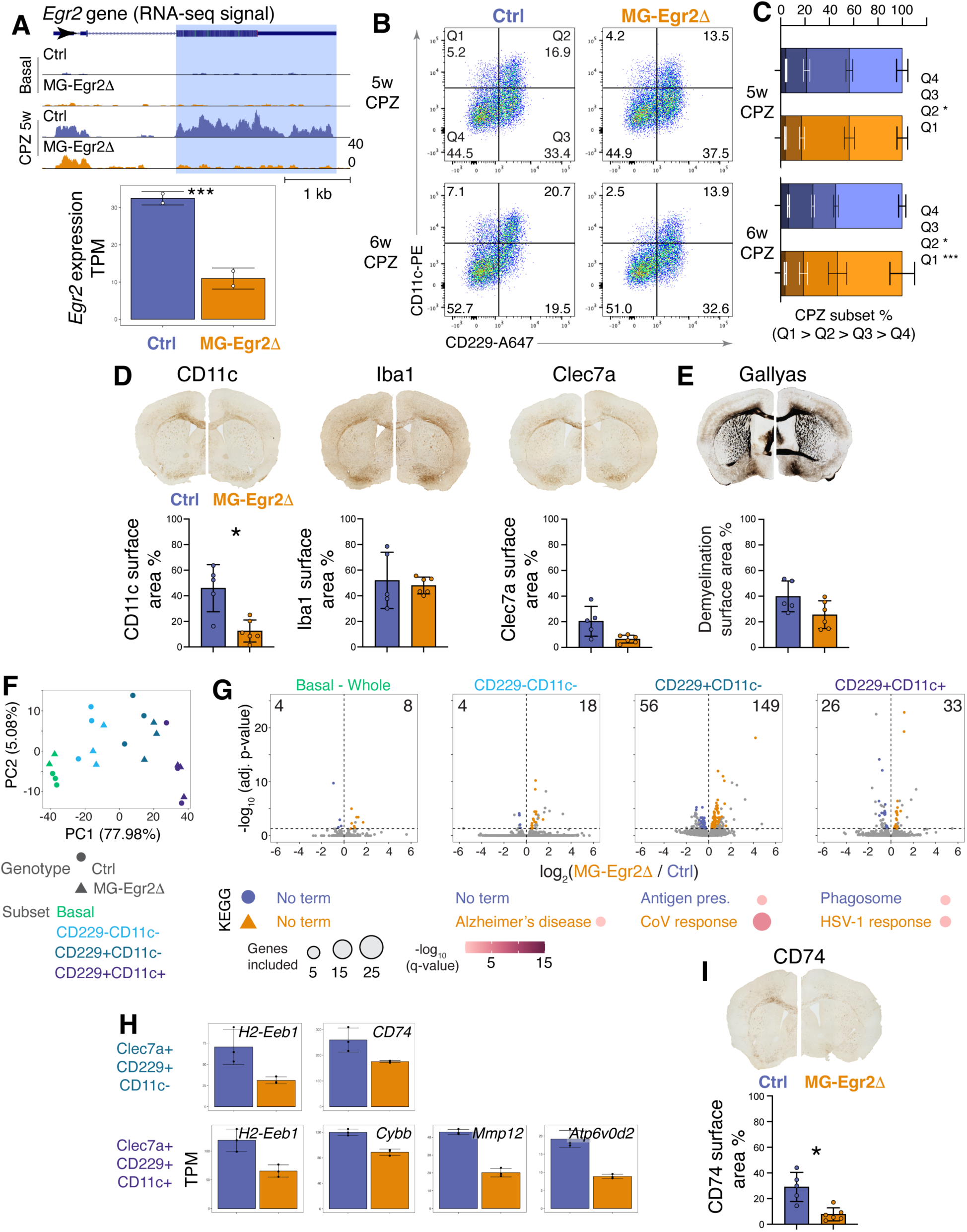
Decreased *Egr2* mRNA expression impairs microglial subsets polarization and lessens myelin damage in mice fed cuprizone diet. **(A)** UCSC genomic browser display of RNA-seq data at *Egr2* locus, and *Egr2* reads (TPM) quantification, comparing microglia from Ctrl and MG-Egr2Δ mice after 5 weeks of CPZ diet. *Egr2* floxed exon is highlighted in blue (Wald test performed with DESeq2 with RNA-seq data; ***q < 0.001; *n* = 2). **(B, C)** Flow cytometric assessment (B) and quantification (C) of CPZ microglia subset, as defined by expression of CD229 and CD11c, isolated from Ctrl and MG-Egr2Δ mice fed CPZ for 5 and 6 weeks (Q1 to Q4) (multivariate ANOVA (MANOVA) followed by a Games-Howell post-hoc test; *n* = 5). **(D, E)** Representative brain microphotographs depicting Iba1, Clec7a, and CD11c immuno-labeling (D), and modified Gallyas staining (E), in Ctrl and MG-Egr2Δ mice fed CPZ for 6 weeks. Below, quantifications of positive (D) or negative (E) area coverage over corpus callosum and external capsule (CD11c, Iba1, Clec7a, Gallyas: normal distribution, unpaired Welch’s t-test, two-tailed; *n* = 5-8). **(F)** Principal component analysis of RNA-seq data from microglia subsets from Ctrl and MG-Egr2Δ mice fed regular or CPZ for 6 weeks (n = 3). **(G)** Volcano plots of RNA-seq data comparing CPZ microglia subsets from Ctrl and MG-Egr2Δ mice. Differentially expressed genes are colored coded (DESeq2, FDR ≤ 0.05; *n* = 3). Below, dominant KEGG term for differentially expressed genes in each subset. **(H)** mRNA expression levels (TPM) of select genes that are expressed at significantly lower levels in MG-Egr2Δ CPZ CD229+CD11c- and CPZ CD229+CD11c+ microglia compared to Ctrl counterparts (linked to G). **(I)** Representative brain microphotographs depicting CD74 immuno-labeling in Ctrl and MG-Egr2Δ mice fed CPZ for 6 weeks. Below, quantification of positive area coverage over corpus callosum and external capsule (normal distribution, unpaired Welch’s t-test, two-tailed; *n* = 5-6). All bar graphs are means ± S.D.; ∗p < 0.05, ∗∗p < 0.01, ∗∗∗p < 0.001.

We next proceeded with RNA-seq and ATAC-seq analyses to unravel how defects in Mef2a activity may disrupt microglia inflammatory polarization and functions to aggravate neural damage in the CPZ model. For this, we sorted 4 subsets, as defined by expression levels of Clec7a, CD229 and CD11c. Principal component analyses and pair-wise comparisons both indicated that the knock-down of Mef2a expression altered gene signatures of all four subsets compared to their matching Ctrl counterparts (Figures 5H, 6I and Table S5). Genes that were consistently upregulated in MG-Mef2aΔ microglia and that could possibly mediate excessive microglial accumulation and damage included protease *Ctse*, and immunoregulatory protein *Isgf8* and *Ifi202b* (Figure S4A). Of note, *Ifi202b*, which possesses apoptosis-inhibitory activity^40^, was never expressed in Ctrl microglia, but was highly induced and expressed in all Mef2aΔ microglia subsets in the demyelinating brain; this may thus contribute to the excessive Iba1+ microgliosis observed in the brain of MG-Mef2aΔ mice fed CPZ. Considering matched pairs individually, CD229+CD11c- microglia from MG-Mef2aΔ mice expressed at higher levels a set of genes involved in “cancer”, which is also coherent with the higher proportion of Ki67+ microglia in these mice (Figure 5I). In contrast, all Clec7a+ subsets from MG-Mef2aΔ mice displayed a consistent defect in induction of phagosomal genes. We also note that several genes typically associated with the DAM signature, including *Cybb*, *CD74*, *H2-Eb1*, *Gpnmb*, *Lgals3*, *Igf1*, as well as *Mmp12* and *Mmp14*, were expressed at lower levels in MG-Mef2aΔ subsets (Figure S4B).

ATAC-seq analyses suggested that the disruption of Mef2a activity negatively impacted the overall signaling configuration in microglia in the demyelinating brain. For example, promoter-distal sites more accessible in either of CD229+CD11c- or CD229+CD11c+ subsets from Ctrl mice encoded motifs linked to C/ebp-AP-1 heterodimer, Irf/Pu.1-Irf, Runx, Egr and Mef2; additional sites in the CD11c+ subset also displayed enrichment for the Atf motifs (Figures 5J and 6K). Of interest, the diminished Egr and Atf epigenomic signatures in CD11c+ Mef2aΔ microglia may be linked to impaired expression of *Egr2* and *Atf5* mRNA relative to the Ctrl subset (Figure 5L). Finally, several TFs involved in cell cycle regulation, including *Etv4*, *Myc*, *Srebf1* and *E2f3*, were expressed at higher levels in Mef2aΔ CD229+CD11c- and CD11c+ subsets, further underscoring the enhanced proliferative activity of microglia with impaired Mef2a activity (Figure S4C).

Collectively, these data indicate that Mef2a provides non-redundant input in microglia that assists in coordinating their proliferative activity and inflammatory state polarization in context of complex brain lesions. In particular, it may critically calibrate the overall cell fitness parameter of the inflammatory microglia population, which in turn could directly impact its net outcome on tissue-remodeling and damage in lesioned context.

### Egr2 promotes CD11c+ polarization and deployment of tissue-remodeling and antigen-presentation gene programs

Finally, we next tested the hypothesis that Egr2 activity is necessary to specify inflammatory state and functions of microglia in the demyelinating brain. Using similar inducible gene deletion strategy as above, we generated Cx3cr1^CreERT2/WT^:Egr2^fl/fl^, henceforth referred to as MG-Egr2Δ and Cx3cr1^CreERT2/WT^, i.e., Ctrls. Based on RNA-seq data, this approach decreased by ∼65% *Egr2* mRNA expression in whole microglial population in the brain of mice fed CPZ for 5 weeks (Figure 6A and Table S6). While flow cytometry data revealed a relatively mild effect on abundance of microglial subsets at CPZ-5w, inhibition of *Egr2* mRNA expression compromised more severely the proportion of CD229+CD11c+ cells at the 6-week CPZ time point as well as expression CD11c on the global microglial cell population (Figures 6B and 7C). Microscopy analyses also confirmed a marked decrease in CD11c+ immunostaining signal coverage in the white matter of MG-Egr2Δ mice compared to Ctrls (Figures 6D). Signals for Iba1 and Clec7a were not significantly different (Figure 6D), and disrupting Egr2 activity in microglia did not impact demyelination in the white matter brain regions of (Figure 6E).

To understand how Egr2 activity may regulate microglial subset polarization and activity, we compared microglia gene expression and genome-wide chromatin accessibility patterns from MG-Egr2Δ and Ctrl mice. First, interfering with Egr2 activity had a relatively minor effect on the number of genes differentially expressed between each subset pairs (Figures 6F, 7G and Table S6). The most pronounced, albeit still relatively mild, effect occurred with the CD229+CD11c- subset, with inhibition of Egr2 activity impairing expression of 56 genes but promoting that of 149; KEGG analysis established a weak overlap between a subgroup of genes among these 149 and antiviral immunity. Regarding the CD229+CD11c+ subset, 26 and 33 genes had respectively lower and higher expression in MG-Egr2Δ mice. However, no subgroup among these differentially expressed genes overlapped with robust q-values with functional KEGG pathways. That said, parsing through the list of transcripts expressed at lower levels in MG-Egr2Δ mice fed CPZ revealed that the full induction of several potent mediators of inflammatory and tissue-remodeling functions critically depends on efficient Egr2 activity (Figure 6H). These comprised matrix metallopeptidase *Mmp12*, superoxide-producing enzyme *Cybb*/Nox2, lysosomal proton pump *Atp6v0d2*, and effector of antigen presentation *H2-Eb1*. Transcripts for *H2-Eb1 mRNA*, as well as *CD74*, were also less expressed in MG-Egr2Δ vs Ctrl CD229+CD11c- microglia; immunostaining analyses confirmed a significantly lower CD74+ signal in the white matter of MG-Egr2Δ mice at the CPZ-6w time point (Figures 6I).

Collectively, these data support a key role for Egr2 in coordinating microglial inflammatory subset activity in the demyelinating brain.

## Discussion

Here, we demonstrate that in the context of brain demyelination, microglia achieve distinct inflammatory states and functions through the concerted activity of different combinations of transcription factors. These include pan-acting regulators of macrophage ontogeny and functions, like Pu.1, AP-1, Cebp and Runx, as well as more context-selective factors such as Mef2, Egr, Bhlhe40, Atf, Irf, Nf-κB and E-box binding family members. Notably, activity of several factors depends on their own transcriptional induction, making the transcriptional process itself a critical gatekeeping mechanism for the coordination of state polarization. Mechanistically, this gatekeeping mechanism implicates delivery of pro-transcriptional input, as well as disengagement of epigenetic-mediated inhibition of transcription, in a locus-dependent manner. Ultimately, the efficient interplay between TFs and the epigenome reprograms the transcriptional output of microglia, enabling these cells to deploy an elaborate spectrum of inflammatory, tissue-remodeling capabilities.

Our analyses revealed that an array of microglia inflammatory states defined by distinct expression patterns of Clec7a, CD229 and CD11c proteins emerge in the demyelinating brain. These results are highly concordant with a recent study that also identified Clec7a+CD11c- and Clec7a+CD11c+ inflammatory microglia in the brain of 5xFAD mice, a model of AD^41^. In that study, the Clec7a+CD11c+ state was exclusively found in close spatial proximity to amyloid deposits, while Clec7a+CD11c- microglia were more broadly distributed. These observations, combined with our results, suggest that these states are intrinsic to microglial inflammatory response and pervasive across chronic brain pathologies, at least in mice. Furthermore, while these inflammatory subsets likely share similar inflammatory capabilities towards endocytic and phago-lysosomal functions, they also likely contribute distinctively to the inflammatory response at the tissue-level. Based on our results, the inflammatory Clec7a+CD229+CD11c- state is compatible with cell proliferation, and through its high expression of *Fn1* and *Vegfa* it may assist in the assembly of cellular and matrix structures to support brain repair and functions^42,43^. Alternatively, the CD229+CD11c+ state is likely endowed with potent pro-inflammatory, tissue-degrading capabilities, owing to its highest, and sometime exclusive mRNA upregulation of matrix metallopeptidase *Mmp12*, superoxide-producing enzyme *Cybb*/Nox2 and pro-inflammatory cytokine *Il1b*. Lastly, we note that while transcripts for *Clec7a* and *Itgax*/CD11c, are frequently listed together as part of the DAM/MGnD gene signature^5,6,32^, their respective protein expression is ultimately independent of one another. In this regard, our data suggest that Clec7a protein expression may characterize a general state of DAM/MGnD microglia, while the CD11c+ state may represent an end-stage differentiation of the DAM/MGnD state.

Distinct lines of evidence suggest that the CD229+CD11c- phenotype arises as an early stage of pro-inflammatory, heightened immune state of microglia associated with brain lesions. First, the downregulation of homeostatic input provided by Mef2a was sufficient to initiate a CD229+ polarization state associated with increased expression of *Apoe*, *Axl* and *Ctse* transcripts, among others. Second, CD229+CD11c- microglia accumulated at CPZ-induced demyelinating lesions in Trem2^-/-^ mice, but these failed to induce expression of *Atf3*, *Bhlhe40* and *Egr2* that may trigger and/or sustain the CD11c+ state. In both contexts, essentially no CD11c+ microglia exist within the larger CD229+ subset. Exactly how the CD229+CD11c- inflammatory state is initiated remains to be determined. One hypothesis is that dampening in inhibitory, homeostatic input could elevate relative strength of pro-inflammatory input provided by potent immune transcription factors such as Pu.1 and Irf8 that specify microglial cell identity in the healthy brain. Our results on Mef2a could support this scenario, as do studies that showed enhanced inflammatory features of microglia in absence of lesions following loss-of function of Mef2c, component of the transforming growth factor receptor - Smads axis and the transcriptional co-repressor Sall1^32,39,44-46^. Lastly, the upregulation of CD229 may be functionally relevant to an expanding microglial population. Indeed, CD229 receptor activation provides a “don’t eat me” signal^47^; it’s early upregulation may be necessary to promote and sustain microglial expansion and minimize probabilities of the microglia population cannibalizing itself.

While perhaps involved in initiating early inflammatory activity, the regulatory mechanisms described above is not sufficient to promote the full spectrum of microglial inflammatory states and functions. This requires both Trem2-independent and Trem2-dependent pro-activating signals to come into play. Our data suggest that the former can mediate early lesion microglia cell proliferation and partial CD229+CD11c- differentiation. On a qualitative level, Trem2-independent signals may coordinate activity of Pu.1, AP-1, C/ebp, Runx, Irf and subset of E-box-binding factors like Hif1α. However, achieving complete CD229+CD11c- state polarization and differentiation into CD229+CD11c+ clearly requires Trem2-dependent signals. Here, our data directly connects Trem2-dependent signaling to de novo, lesioned-induced activity of Atf3, Bhlhe40 and Egr2 transcription factors, among others. In turn, their joint activity may promote the emergence and/or sustaining of a CD11c+ state that is endowed with potent tissue-remodeling and potentially antigen-presenting functions, as suggested by Egr2 loss-of-functions experiments. A link between Egr2 input in supporting high tissue-remodeling capabilities is also corroborated by previous studies that suggested that Egr2 underlies alveolar macrophages activity; these are indeed highly efficient tissue-remodelers owing to their roles in pulmonary surfactant degradation and residence in complex, challenging immune environment^48,49^. That said, while Trem2 signaling promotes the induction of Egr2, Atf3 and Bhlh40, which a prerequisite for their activity, the extent to which it also directly controls their transcriptional activity at the protein level remains to be established.

In sum, our study clarifies our understanding of the signaling, epigenomic and transcriptional mechanisms that underlie microglial inflammatory state polarization and functions. Given the intricate association between microglial inflammatory activity and neurodegenerative disorders, these new insights may inform on the dominant coordinators of microglial roles in these disorders and may thus assist in the development of more efficient therapeutics.

## Resource availability

### Lead contact

Further information and requests for resources and reagents should be directed to and will be fulfilled by the lead contact, David Gosselin.

### Materials availability

All requests for resources and reagents should be directed to and will be fulfilled by the lead contact. This study did not generate new unique reagents.

### Data and code availability

RNA-seq and ATAC-seq will be made publicly available publication in a peer-reviewed journal. This study did not generate new code. Any additional information required to reanalyze the data reported in this paper is available from the lead contact upon request.

## Acknowledgements

The authors thank Sophie Vachon, Stephanie Bernard and Emilie Méthot-Lauzé for assistance with mouse husbandry. This study was supported by the following grant allocated to David Gosselin: Hélène-Hallé new investigator grant, NARSAD new investigator award (no. 27359), Scottish Rite Charitable Foundation of Canada, Fondation du CHU de Québec new investigator grant, Faculté de Médecine de l’Université Laval startup funds, Centre de Recherche du CHU de Québec startup funds, and additional support from Axe Neuroscience du CRCHU-CHUL. These studies were also supported by a Canadian Institutes of Health Research project grant, MS Canada discovery grant and Alzheimer’s Society of Canada new investigator biomedical grant allocated to David Gosselin, and a Foundation grant from the Canadian Institutes of Health Research allocated to Serge Rivest. Félix Distéfano-Gagné currently holds a Doctoral Research Award from the Canadian Institutes of Health Research and previously held Master’s awards from the Canadian Institutes of Health Research and from Fonds de Recherche du Québec – Santé. Serge Rivest holds a Canada Research Chair in Neuroimmunology. David Gosselin is supported by a J2 award from Fonds de Recherche du Québec – Santé. This publication includes data generated at the UC San Diego IGM Genomics Center utilizing an Illumina NovaSeq X Plus that was purchased with funding from a National Institutes of Health SIG grant (#S10 OD026929).

## Author contributions

FDG, NB, AMX and DG conceived the study. FDG, NB, SF and FDG performed mice experiments, collected and analyzed data. WS and SB collected data. SR provided intellectual input. DG generated sequencing libraries and analyzed data. FDG, NB and DG wrote the manuscript with assistance from AMX. DG supervised all aspects of the project.

## Methods

### Mice

C57BL/6J (The Jackson Laboratory, strain 000664), C57BL/6J-Trem2^em2Adiuj/J^ (Trem2^-/-^; The Jackson Laboratory, strain 027197) and B6.129P2(C)-Cx3cr1^tm2.1(cre/ERT2)Jung/J^ (Cx3cr1^CreERT2^; The Jackson Laboratory, strain 020940) were first purchased from The Jackson Laboratory. Mef2a^fl/fl^ and Egr2^fl/fl^ mice were generously donated, respectively, by Dr. Eric Olson (University of Texas Southwestern), and Dr. Warren Leonard and Dr Vanja Lazarevic (National Institutes of Health). All mice were backcrossed on a C57BL/6 background for at least 5 generations. Cx3cr1^CreERT2^ mice were bred with Mef2a^fl/fl^ or Egr2^fl/fl^ to create Cx3cr1^CreERT2/WT^::Mef2a^fl/fl^ (MG-Mef2aΔ) or Cx3cr1^CreERT2/WT^::Egr2^fl/fl^ (MG-Egr2Δ) mouse lines. Cx3cr1^CreERT2/WT^ mice were used as controls (Ctrls). Internal colonies for each of these mouse lines were established upon reception of breeding pairs at the animal research facility of the Centre de Recherche du Centre Hospitalier Universitaire de Québec – Université Laval. Males were used throughout this study. All adult mice were of 3 to 4.5 months of age at sacrifice. Mice had ad libitum access to food and water. All animal procedures were approved by Université Laval’s animal care and ethics committee (CPAUL3) and performed in compliance with ethical regulations and guidelines of the Canadian Council on Animal Care. For sampling, mice from at least 2 litters were included in each group, and one litter could not contribute more than 3 mice.

### Tamoxifen administration

Tamoxifen (Sigma-Aldrich, T5648) was dissolved in 10% ethanol (Thermo Fisher, A962P-4) and 90% corn oil (Sigma-Aldrich, C8267). MG-Mef2aΔ and MG-Egr2Δ mice were administered daily intraperitoneal injections (100 mg/kg) over 4 consecutive days starting between postnatal days 35 to 42 to induce deletion of floxed alleles by the CreERT2 recombinase^37,38^. Mice were then left undisturbed for 6 to 8 weeks after the last injection. Controls (Ctrls) underwent the same procedure.

### Cuprizone diet

Cuprizone (Sigma-Aldrich, 14690) was mixed with regular ground chow (0.2% wt/wt) and fed ab libitum to mice for 4, 5 or 6 weeks^50^. Diet was changed every 2 days.

### Method Details

#### Flow cytometry and fluorescence-activated cell sorting (FACS)

Mice were euthanized with an overdose of ketamine-xylazine solution (90 ml/kg & 10 ml/kg) injected intraperitoneally and subsequently perfused transcardially with ice-cold 0.9% NaCl saline solution (Sigma-Aldrich, S5886-1K6). Brains were extracted and placed in ice-cold base buffer (BB) solution (HBSS 1X (Life Technologies, 14175-095), 1% BSA (Sigma-Aldrich, A9647), 1mM EDTA (Thermo Fisher Scientific, 15575020), 1 mM sodium butyrate (Sigma-Aldrich, B5887), 1 mM Flavopiridol (Sigma-Aldrich, F3055))^35^. For flow cytometry, brains were cut into smaller pieces in BB solution before adding Accutase solution (Sigma-Aldrich, A6964) to enzymatically dissociate the brain. After 1 hour of incubation, brains were further homogenized on ice using a cut P1000 tip, then an uncut tip and finally a P200 tip. For FACS, brains from 2-6 mice were from same conditions were pooled and homogenized in BB solution by gentle mechanical dissociation on ice using a 7 ml polytetrafluoroethylene pestle (Duran Wheaton Kimble, 357542) and filtered through a 100 µm cell strainer (ThermoFisher 22-363-549).

Following brain homogenization, homogenates were resuspended in 37% Percoll (Thermo Fisher Scientific (Cytivia) 17089101; 8 ml/brain, in 15 ml tubes (1 tube/brain), which were then centrifuged at 600 x g for 20 min at 20°C, with medium acceleration and no deceleration. Pellets were then recovered and washed once in with BB solution, centrifuged at 300 RCF at 4°C for 7 min, resuspended in 4 ml HBSS 1X and then centrifuged at 300 RCF at 4°C for 7 min to remove proteins. Cells were resuspended in 100 µl HBSS1x and incubated on ice in the dark for 10 min with 1:100 anti-CD16/CD32 receptor unconjugated antibody (BioLegend 101302, clone 93). Then, cells were incubated on ice in the dark for 10 min with 1:1000 Live/Dead yellow stain (Thermo Fisher Scientific L34959) and conjugated antibodies directed against cell surface proteins: 1:167 anti-Ace-Alexa 750 (R&D Systems FAB15131S, clone 230214), 1:100 anti-CD11b-Brilliant Violet 421 (BioLegend 101251, clone M1/70), 1:100 anti-CD44-APC/Cy7 (BioLegend 103027, clone IM7), 1:200 anti-CD45-BB700 (BD Biosciences 566439, clone 30-F11), 1:100 anti-CD229-Alexa 647 (BD Biosciences 566689, clone Ly9.7.144), 1:133 anti-CD11c-PE (BioLegend 117308, clone N418) and 1:100 anti-Clec7a-PE/Cy7 (Thermo Fisher Scientific 25-5859-80, clone bg1fpj). After incubation, cells were washed in HBSS1x and centrifuged at 300 RCF at 4°C for 7 min. Then, cell pellets were either 1) resuspended in BB and filtered through a 35 µm cell strainer and analyzed or sorted for RNA-seq and ATAC samples, or 2) underwent intracellular staining for Ki67 for analysis. For Ki67 staining, cell pellets were resuspended in HBSS 1x and fixed with 1% PFA at room temperature (RT) in the dark for 10 min, then permeabilized with eBioscience Foxp3 / Transcription Factor Fixation/Permeabilization Concentrate and Diluent (Thermo Fisher Scientific 00-5521-00) and incubated with 1:100 anti-Ki67-Alexa 488 antibody (Thermo Fisher Scientific, 53-5698-82, clone SolA15) on ice in the dark for 30 min. After incubation, cells were either washed in BB, centrifuged at 1000 RCF at 4°C for 10 min and filtered through a 35 µm cell strainer using BB and then analyzed. For cross-linking, cell pellets were resuspended in HBSS 1x and fixed with 1% PFA at RT in the dark for 10 min, after which 1/20^th^ volume of 2.625 M glycine (Thermo Fisher Scientific BP381-500) was added for quenching for 10 min. Cells were washed in BB, centrifuged at 1000 RCF at 4°C for 10 min and filtered through a 35 µm cell strainer using BB and then sorted. Flow cytometry data was acquired with a BD FACSymphonyA1 instrument. Data was analyzed with FlowJo (v10.10.0). Microglia were sorted with BD FACSAria Fusion instrument (100 µm nozzle).

#### Immunohistochemical staining and analysis

For histological analyses, transcardial perfusion with 0.9% NaCl was immediately followed perfusion with fixative solution (4% paraformaldehyde (Electron Microscopy Sciences 19202) in 0.1M Borax buffer (pH 9.0)). Brains were extracted and placed in same fixative solution for two days for post-fixation, after which they were transferred to a mix of fixative solution containing 20% sucrose. Brains were cut the next day at a thickness of 30 µm by Leica SM2010 microtome, and sections were stored in the anti-freeze solution (0.006% NaH2PO4 (Thermo Fisher Scientific BP329-500), 0.0205% Na2HPO4 (Thermo Fisher Scientific BP332-500), 30% ethylene glycol (Sigma-Aldrich 102466-5L) and 20% glycerol (Thermo Fisher Scientific BP229-4), in water) at –20°C.

Brain sections were rinsed 3 times for 10 min in KPBS (21.87mM K2HPO4 (Thermo Fisher Scientific BP363-1), 3.3 mM KH2PO4 (Thermo Fisher Scientific BP362-500), 138.6 mM NaCl, in water, pH adjusted to 7.2 with HCl) before incubation for 10 min in 3% H2O2 (Sigma-Aldrich 323382). After one KBPS wash, sections were incubated in blocking solution (1% Triton-X-100, 4% goat serum, 1% BSA, in KPBS) for 1 hour. Unconjugated primary antibody diluted in blocking solution took place overnight, using one of the following antibodies: 1:1500 rabbit anti-CD11c (New England Biolabs 97585, clone D1V9Y), 1:2000 rat anti-CD74 (BD Biosciences 555317, clone In-1), 1:500 rat anti Clec7a (Invivogen mabg-mdect, clone R1-8g7), or 1:1250 rabbit anti-Iba1 (Wako Fujifilm 019-19741). The next morning, sections were rinsed 3 times with KPBS and incubated in diluted biotinylated secondary antibody (either 1:1500 goat anti-rabbit-biotin (Vector Labs, BA-1000) or 1:1000 goat anti-rat-biotin (Thermo Fisher Scientific 31830), in KPBS) for 2 hours. After 3 rinses, slices were incubated for 1 h in a Vectastain Elite ABC-HRP Kit (Vector Laboratories PK-6100) solution (2.25 µl of each reagent per ml of KPBS) to add peroxidase. Rinses took place before revelation in a 3,3’-diaminobenzidine tetrahydrochloride hydrate (DAB; Sigma-Aldrich 06522) solution (0.5 mg/ml DAB, 0.003% H2O2, in KPBS) for 7 to 10 minutes. After rinsing, brain slices were mounted on positively charged slides (VWR Microslides 48311-703), dehydrated in EtOH baths, rehydrated in xylene and coverslipped with DPX mountant (Sigma-Aldrich 06522).

Positive signal area coverage for Iba1, Clec7a, CD11c and CD74 over white matters regions, including corpus callosum and adjacent external capsule (see Figure S1B), were analyzed. For this, 2 to 3 microphotographs of brain slices between Bregma 0.13 mm and 1.09 mm, as defined by Paxinos and Franklin’s mouse brain stereotaxic coordinates (4^th^ edition), were taken using an Olympus BX50 microscope mounted with a QImaging Retiga EXi Fast CCD digital camera. Images were captured in QCapture version 2.9.10 and positive staining area was quantified over white matter regions using the Pixel Classifier tool in QuPath version 0.5.1^51^. Representative slices were rephotographed at higher magnification and stitched using Fiji’s Grid/Collection stitching plugin^52^.

For quantification of Iba1+ microglia cell density, 2 microphotographs centered on the primary somatosensory cortex were taken on adjacent brain slices, between Bregma 0.13 mm and 1.09 mm, and then imported to ImageJ^53^. A 0.73 mm x 0.73 mm box was first drawn over primary somatosensory cortex and encompassed Iba1^+^ cells were then counted. Cell counts over the two slices for each region were then averaged.

All analyses were performed blindly.

#### Gallyas myelin staining and analysis

To stain myelinated axons, we adapted the Gallyas silver stain technique, in which colloidal silver binds myelin^54,55^. Brain sections were first mounted on positively charged slides in KPBS and allowed to dry for 2 days under vacuum. Slides were then rinsed in KPBS before incubation in a 2:1 pyridine (Thermo Fisher Scientific P368) and acetic anhydride (Sigma-Aldrich 320102) solution for 1 hour. They were then successively rinsed in 50% and 30% EtOH, then 0.05% and 0.1% acetic acid (Sigma-Aldrich,695092) before 3 rinses in 0.5% acetic acid. Slides were then incubated for 45 minutes in a basic silver nitrate solution (0.2% ammonium nitrate (Sigma-Aldrich A9642), 0.22% silver nitrate (Thermo Fisher Scientific S181-25), 6 mM NaOH (Sigma-Aldrich S5881-500G), in water). After three rinses in 0.5% acetic acid, they were physically developed in an acidic silver nitrate and sodium carbonate solution (2.5% sodium carbonate anhydrous (Thermo Fisher Scientific S263), 0.1% ammonium nitrate, 0.1% silver nitrate, 0.5% tungstosilicic acid hydrate (Sigma-Aldrich T2786), 0.0146% PFA pH 7.4 (Thermo Fisher Scientific J61899-AK), in water) for 30 to 45 min. A single rinse in 0.5% acetic acid took place before quick incubations in 0.2% potassium ferricyanide (Sigma-Aldrich 393517) for 10 to 30 seconds and 0.5% sodium thiosulfate (Sigma-Aldrich S1648) for 2 minutes. Finally, slides were rinsed three times in KPBS.

Coverslip and Gallyas image acquisition over white matters regions was performed as described above for immunohistochemical stains of Iba1, Clec7a, CD11c and CD74. To infer damage, negative area signal coverage values were quantified in QuPath using the Pixel Classifier tool, as opposed to positive signal coverage values. All analyses were performed blindly.

#### Immunofluorescence staining and analysis

Brain sections were rinsed 3 times in KPBS before incubation in blocking solution (1% Triton-X-100, 4% donkey serum, 1% BSA, in KPBS) for 1 hour and overnight incubation with the following unconjugated primary antibodies: 1:500 goat anti-Iba1 (Sigma-Aldrich SAB2500041), 1:500 rat anti-Clec7a (Invivogen mabg-mdect, clone R1-8g7) and 1:1000 rabbit anti-Cd11c (New England Biolabs 97585S, clone D1V9Y), in blocking solution. Afterwards, they were rinsed and incubated in the dark with the following secondary antibodies: 1:1000 donkey anti-goat-Alexa 647 (Thermo Fisher Scientific A21447), 1:1000 donkey anti-rat-Alexa 555 (Thermo Fisher Scientific A48270) and 1:1000 donkey anti-rabbit-Alexa 488 (Thermo Fisher Scientific A32790), in KPBS. After rinsing, sections were mounted on positively charged slides and coverslipped using Fluoromount-G Mounting Medium (Thermo Fisher Scientific 00-4958-02).

Z-stacked and tiled microphotographs were taken using a Zeiss LSM 800 confocal microscope equipped with a Zeiss Axiocam 305 color camera. Images were captured and constructed using Zeiss Zen version 2.6.

#### RNA-seq library preparation: Isolation and fragmentation of poly(A) and cDNA synthesis

RNA-seq library preparation was performed similarly as previously described^35^. Following sorting, 100,000 to 200,000 microglia were centrifuged (350 RCF at 4°C for 10 min), and resuspended into 100 µl lysis/Oligo d(T) Magnetic Beads binding buffer (Thermo Fisher Scientific A33562, or 100 mM Tris-HCl pH 7.5, 500 mM LiCl, 10 mM EDTA pH 8.0, 1% LiDS, 5 mM DTT, in molecular grade water) and stored at -80°C until processing. For isolation and enrichment of polyA mRNA, samples were thawed at RT and incubated with 20 µl Oligo d(T) Magnetic Beads (NEB S1419S) as follows: 2 min at 65°C on a PCR cycler and then 10 min at room temperature (RT). Samples were then placed on a collection magnet at RT and washed serially with 180 µl of washing buffer 1 (RNA-WB1; 10 mM Tris-HCl pH 7.5, 0.15 M LiCl, 1 mM EDTA pH 8.0, 0.1% LiDS, 0.1% Triton X-100, in molecular grade water) and then washing buffer 3 (RNA-WB3; 10 mM Tris-HCl, 0.15 M NaCl, 1 mM EDTA pH 8.0, in molecular grade water). RNA was eluted from Oligo (t) beads by resuspending the beads in 50 µl of elution buffer (EB; 10 mM tris-HCl pH 7.5, 1 mM EDTA pH 8.0, in water) followed by incubation at 80°C for 2 min on a PCR cycler. The samples were then placed back on the magnet, and the elution buffer supernatant containing the poly(A) RNA was carefully collected and placed on ice.

A second round of poly(A) RNA selection was then performed. Eluted Oligo d(T) beads were first washed on a magnet with 150 µl EB and then 150 µl 2X Oligo d(T) binding buffer (2x DTBB; 20 mM Tris-HCl pH 7.5, 1 M LiCl, 2 mM EDTA pH 8.0, 1% LiDS, 0.1% Triton X-100, in molecular grade water) and resuspended in 50 µl of 2x DTBB. Resuspended Oligo d(T) beads were then added to isolated RNA and incubated at 65°C and at RT as described above. Beads were then placed on a magnet and washed once with 180 µl of RNA-WB1, then once with 180 µl of RNA-WB3, and then resuspended in 30 µl of 1x First-Strand buffer (made from 5x stock Thermo Fisher Scientific 18080-044 with molecular grade water) transferred in a new PCR strip. Beads were collected on a magnet and resuspended in 10 µl of 2x SuperScript III buffer (made from 5x stock Thermo Fisher Scientific 18080-044 supplemented with 10 mM DTT, in molecular grade water). Resuspended beads were then incubated at 94°C for 9 min to fragment poly(A) RNA. Beads were then placed back on a magnet, and the supernatant containing poly(A) RNA was carefully collected and placed on ice.

Collected poly(A) mRNA in 2X SuperScript III buffer was then incubated for 1 min at 50°C on a PCR cycler with 2.5 µl of the following mix: 1.5 µg Random Primer (Thermo Fisher Scientific 48190-011), 10 µM Oligo d(T)20 primer (Thermo Fisher Scientific 18418020), 10 units SUPERase•In RNase Inhibitor (Thermo Fisher Scientific A32694), 4.0 mM dNTP mix (Thermo Fisher Scientific 18427088). Samples were then immediately placed on ice for 5 min. First-strand synthesis was then performed by incubation at 25°C for 10 min and 50°C for 50 min on a PCR cycler with 7.6 µl of the following mix: 0.2 µg actinomycin D (Sigma-Aldrich A1410; stock at 2 µg/µl in DMSO), 13.15 mM DTT (Thermo Fisher Scientific 361045836), 0.026% Tween-20 (from 10% stock Enzo 80-1929, in molecular grade water), 100 units SuperScript III Reverse Transcriptase (Thermo Fisher Scientific 18080-044), in molecular grade water. After incubation, RNA/DNA complexes were isolated by adding 36 µl of RNAClean XP beads (Beckman Coulter A63987) and incubated for 10 min at RT and then 10 min on ice. Samples were then placed on a magnet and beads were washed twice with 150 µl of 75% EtOH. Following washings, beads were air-dried at RT for 6-10 min and eluted with 10 µl of water.

Second-strand synthesis was then performed. RNA/DNA samples, in 10 µl of water, were incubated for 2.5 hours at 16°C with 5 µl of the following mix: 3x NEB Buffer 2 (10x stock New England Biolabs B7002S), 1.0 µl PCR mix (stock of 10 mM solution for each dATP, dCTP, dGTP, and dUTP from Thermo Fisher Scientific, in molecular grade water), 2.0 mM dUTP (Thermo Fisher Scientific, R0133), 1 unit RNAse H (Enzymatics Y9220L), 10 units DNA Polymerase I (Enzymatics P7050L), 0.03% Tween-20, in molecular grade water). DNA was then purified by addition of 1.5 µl Sera-Mag SpeedBeads Carboxyl Magnetic Beads (Thermo Fisher Scientific (Cytivia) 45152105050250), resuspended in 30 µl 20% PEG 8000/2.5 M NaCl (PEG 8000 from Sigma-Aldrich P2139; 5M NaCl from Thermo Fisher Scientific AM9759), incubated at RT for 15 min and placed on a magnet for two rounds of bead washing with 80% EtOH. Beads were then air dried for 6-10 min and DNA was eluted from the beads by adding 40 µl of water. Supernatant was then collected on a magnet and placed on ice or stored at -20°C until DNA blunting, poly(A)-tailing, and adapter ligation (see below).

#### RNA-seq final library preparation

Sequencing libraries were prepared from generated cDNA (RNA) by blunting, A-tailing, and adapter ligation as previously described using barcoded adapters (NEXTflex Unique Dual Index Barcodes (Set B), Bioo Scientific NOVA-514151)^56,57^. Prior to final PCR amplification, RNA-seq libraries were digested by 30 min of incubation at 37°C with Uracil DNA Glycosylase (final concentration of 0.134 units per µl of library volume; UDG, Enzymatics G5010L) to generate strand-specific libraries. Libraries were PCR-amplified for 14 cycles and size selected for fragments (200-500 bp for RNA-seq) by gel extraction (10% TBE gels, Thermo Fisher Scientific Technologies EC62752BOX). RNA-seq libraries were paired-end sequenced for 100 or 150 cycles on an Illumina NovaSeq X Plus or NovaSeq 6000 (Illumina, San Diego, CA) according to manufacturer’s instruction.

#### Assay for Transposase-Accessible Chromatin-sequencing (ATAC-seq)

Immediately after sorting, aliquot of 80,000 microglia were centrifuged (350 RCF at 4°C for 10 min) and cell pellets were lysed in 50 µl lysis buffer (10 mM Tris-HCl ph 7.5, 10 mM NaCl, 3 mM MgCl2, 0.1% IGEPAL, CA-630, in molecular grade water) on ice and nuclei were pelleted by centrifugation at 500 RCF for 10 min. Nuclei were then gently resuspended on ice in 50 µl transposase reaction mix (1x Tagment DNA buffer (Illumina 15027866), 2.5 µl Tagment DNA enzyme I (Illumina 15027865), in molecular grade water) and incubated at 37°C for 30 min on a PCR cycler. DNA was then purified with Zymo ChIP DNA concentrator columns (Zymo Research D5205) and eluted with 10 µl of elution buffer. DNA was then amplified with PCR mix (1.25 µM for each of i5- and i7- indexed primers, from Mezger et al. 2018^58^, 0.6x SYBR Green I (Thermo Fisher Scientific S7563), 1x NEBNext High-Fidelity 2x PCR MasterMix, (NEB M0541)) for 9 cycles, run on gel for size selection of fragments (160-280 bp) as detailed above, and paired-end sequenced for 100 or 150 cycles on an Illumina NovaSeq X Plus or NovaSeq 6000 (Illumina, San Diego, CA) according to manufacturer’s instruction.

#### Illumina sequencing

Sequencing for RNA-seq and ATAC-seq samples was performed on NovaSeq X Plus at the IGM Genomics Center, University of California, San Diego, La Jolla, CA, USA, or NovaSeq 6000 at The Centre for Applied Genomics (TCAG) at The Hospital for Sick Children, Toronto, ON, Canada.

#### Sequencing data analysis preprocessing

Only R1 sequencing files were used for alignment and analyses, except for two ATAC-seq files for Trem2^-/-^ CPZ 4 weeks CD229-CD11c- and CD229+CD11c- subsets – replicates 3 for which R2 files were used. For all PE 150 sequencing files, 50 bp were first trimmed off the 3’ end of R1 reads using fastp (fastp-0.20.0; command *-t 50*)^59^. For ATAC-seq R1 sequencing files, adapters were also removed off 5’ end of R1 reads using fastp in default mode. Following these steps, and for all other single- or paired-end 100 RNA-seq data, R1 reads FASTQ sequencing files were mapped to the mouse mm10 reference genome. STAR (2.7.0f) with default parameters was used to align RNA-seq experiments to the GRCm38 -mm10 mouse genome^60^. Bowtie2 (2.3.4.1) with default parameters was used to align ATAC-seq experiments to the GRCm38 -mm10 mouse genome^61^. HOMER was used to convert aligned reads into “tag directories” for further analyses^62^.

#### RNA-seq

For differential expression analyses, a table read count including relevant group replicates was first created using the *analyzeRepeats.pl* program of HOMER (v4.11; http://homer.ucsd.edu/homer/) with the following parameters: *rna -noadj - condenseGenes -count exons -pc 3*. The resulting read count dataset was used to produce input files for pairwise comparisons using R package DESeq2^63^. A Wald test was performed with DESeq2 to identify differentially enriched transcripts between groups of interest, using the following parameters: *lfcThreshold = 0*, *alpha = 0.05*. For data plotting and interpretation purposes and functional enrichment analyses with Metascape (see below), only protein-coding mRNAs that have a Transcript Per Million (TPM)-normalized value of 16 or greater in at least one condition were retained. The base-2 logarithm of the TPM values was taken after adding a pseudocount of 1 TPM to each transcript. TPM datasets were generated using HOMER’s *analyzeRepeats.pl* program with parameters *rna -condenseGenes -count exons -pc 3 -tpm*.

For quantification of *Mef2a* floxed exons count (see Figure 4A), a regions.txt file with genomic coordinates including the *Mef2a* floxed exons (chr7:67,295,381-67,295,569) and *Nup98* gene body (chr7:102,119,398-102,210,176) was first generated. The regions.txt file was then annotated with tags from RNA-seq tag directories from homeostatic Ctrl and MG-Mef2Δ whole microglia, using HOMER’s *annotatePeak.pl* program with *mm10 -noadj* parameters. *Mef2a* floxed exons tag counts were then normalized to that of *Nup98* gene body; ratios were then used to assess differential exon expression between genotypes. For *Egr2*, the floxed exon contains the poly(A) sequenced. As such, *Egr2* exon deletion efficient efficiency was assessed as part of mRNA differential analysis above.

Metascape with default parameters was used to perform functional enrichment analyses on groups of genes of interest (RefSeq identifiers). Terms associated with KEGG pathways were used for data interpretation^64,65^.

#### ATAC-seq

Regions with accessible chromatin for each sample were first identified using their tag directory, using HOMER’s *findPeaks.pl* program with the *-style factor* parameter. For group comparison, a table of genomic coordinates containing all the regions identified across the relevant sample replicates was generated using HOMERS’s *mergePeaks.pl* program, with *-size given* parameter. Then, the regions from the mergePeak file were annotated with sequencing tags from ATAC-seq tag directories from relevant sample replicates using HOMER’s *annotatePeak.pl* program with *mm10 -noadj* parameters. Then, regions with differential chromatin accessibility were identified using HOMER’s *getDiffExpression.pl* program, which leveraged R/DESeq2 to assess differential enrichment with the following parameters: *-AvsA -log2fold 0 -fdr 0.05 -simpleNorm*. The resulting file is a read count normalized to 10 million total reads and with differential enrichment results. For data interpretation and plotting, differentially enriched distal CREs, defined as beyond 1000 bp upstream and 500 bp downstream of a TSS, with a LFC of 1 and at least 16 normalized reads were considered.

#### Sequencing data visualization

The UCSC genome browser was used to visualize RNA-seq and ATAC-seq sequencing data^66^. Bedgraph.gz files were generated using HOMER’s *makeUCSCfile.pl*; -strand parameter was specified for RNA-seq files.

#### Quantification and statistical analysis

Data were screened for outliers using Graphpad Prism version 10.4.2’s ROUT function, at a Q value of 1%. Overall, one sample value (Iba1 signal coverage from a WT mouse; see Figure 3E) met qualified as an outlier and was thus removed from statistical analyses and plotting. After normality assessment by the Shapiro-Wilk test, flow cytometry and histological analyses were analyzed using Welsh’s t-test (if normality is not rejected) or the Mann-Whitney test (if normality is rejected) when two groups are compared, or two-way ANOVA followed by Tukey’s multiple comparisons test when more than two groups are compared, using Graphpad Prism version 10.4.2. Differences in population proportions defined by CD229 and CD11c, i.e., quadrants Q1 to Q4, were assessed using a multivariate ANOVA (MANOVA) followed by a Games-Howell post-hoc test. Differential expression analyses were assessed using a Wald test with a false discovery rate of 5% as described above, and subsequent graphs were generated in R version 4.4.1 with the following packages: tidyverse, pheatmap, VennDiagram, rcartocolor, car, and viridisLite. Flow cytometry scatterplots were generated using FlowJo version 10.10.0.

## Supplementary figures

**Supplementary figure 1.**
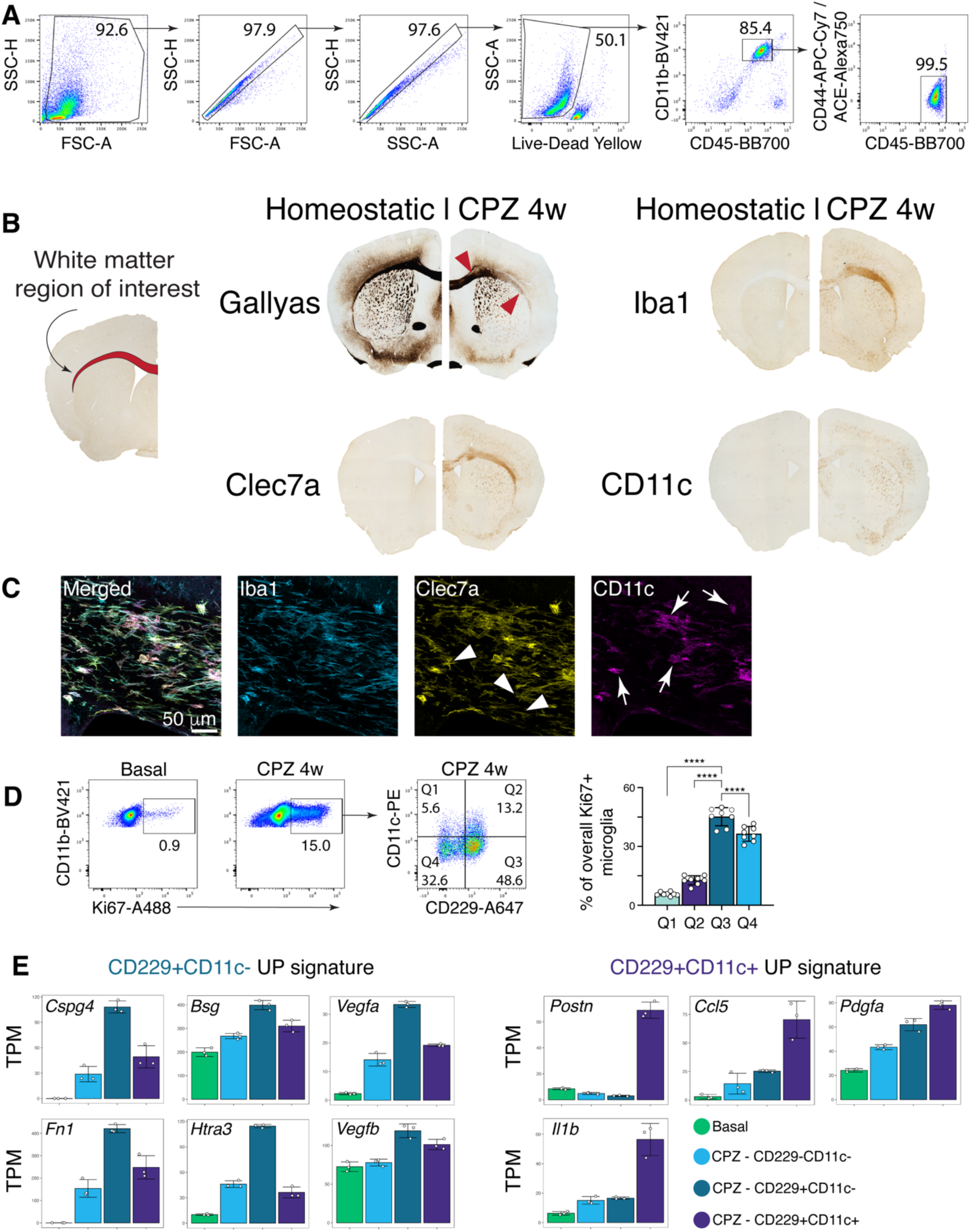
Flow cytometry, microscopy and gene expression data of homeostatic and inflammatory microglia associated with demyelination induced by cuprizone diet, related to figure 1. **(A)** Flow cytometry gating strategy used to define single, live CD11b+CD45^+^ CD44^Low/Neg^Ace^Low/Neg^ microglia. Annotated numbers represent parent gate percentage. **(B)** Representative brain microphotographs of modified Gallyas staining, and Iba1, Clec7a and CD11c immuno-labeling, contrasting signals from brains isolated from mice fed regular diet to mice fed CPZ for 4 weeks. Red arrows point to examples of areas of demyelination. **(C)** Representative confocal microphotograph depicting expression of Clec7a and CD11c on Iba1+ cells in the brain of mice fed CPZ for 4 weeks. Arrowheads depict Iba1+Clec7a+CD11c- cells, while arrows depict Iba1+Clec7a+CD11c+ cells. **(D)** Flow cytometric assessment and quantification of distribution of Ki67+ microglia among the CPZ microglia subsets defined by CD229 and CD11c expression (normal distribution, one-way ANOVA followed be Tukey’s multiple comparison test). **(E)** Bar graphs depicting expression of select genes with significantly higher expression in CPZ CD229+CD11c- vs CD229+CD11c+ subsets, and vice versa (Transcripts Per Million (TPM) normalization). All bar graphs are means ± S.D.; ∗∗∗∗p < 0.0001.

**Supplementary figure 2.**
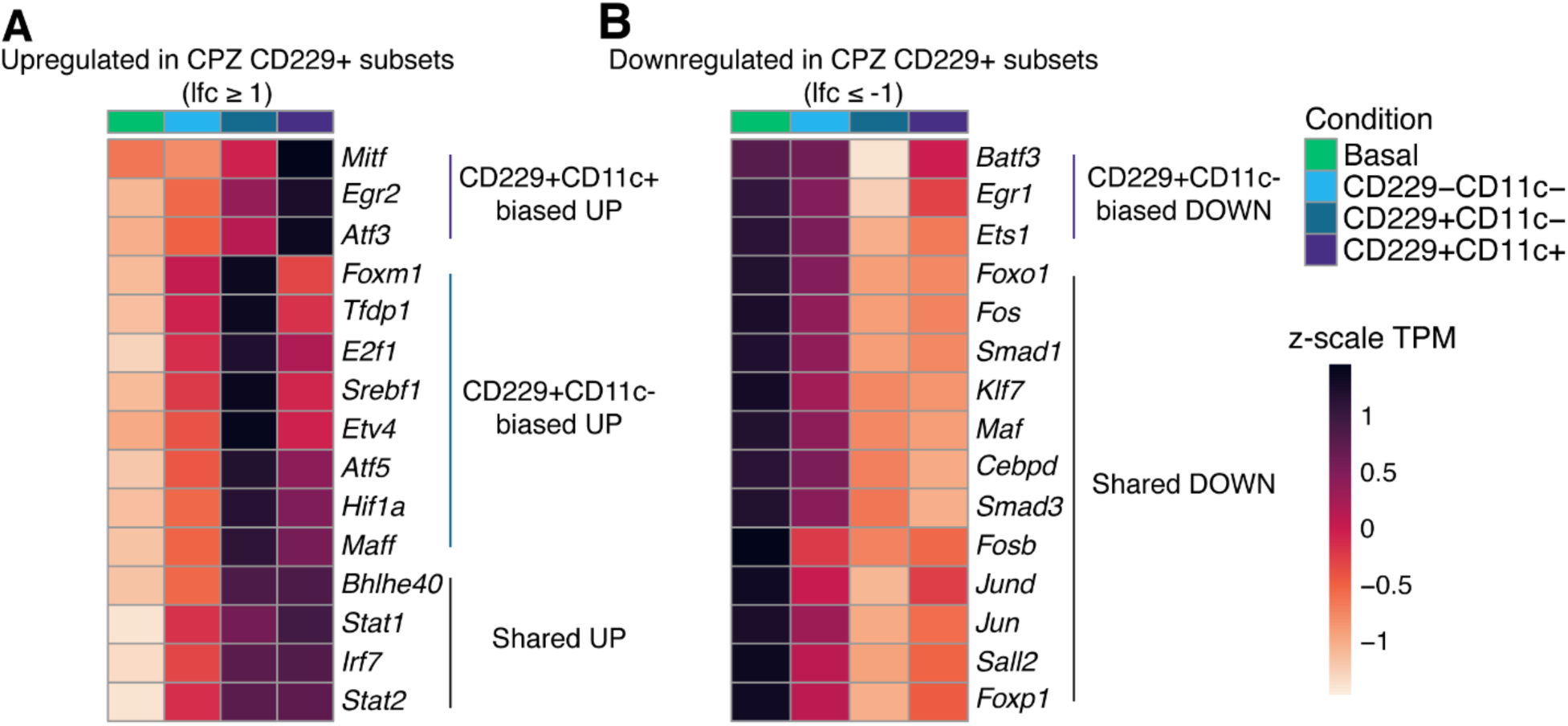
Transcription factor mRNA expression changes in inflammatory microglia subsets compared to homeostatic microglia, related to figure 2. **(A, B)** Heatmaps depicting z-scores of mRNA expression, for significantly modulated transcription factor genes in WT CPZ CD229+ microglia comparing homeostatic and CPZ subset isolated from WT mice. Differentially expressed genes assessed with DESeq2 (FDR ≤ 0.05, lfc ≥ 1 (A) or lfc ≤ -1 (B); n = 3).

**Supplementary Figure 3.**
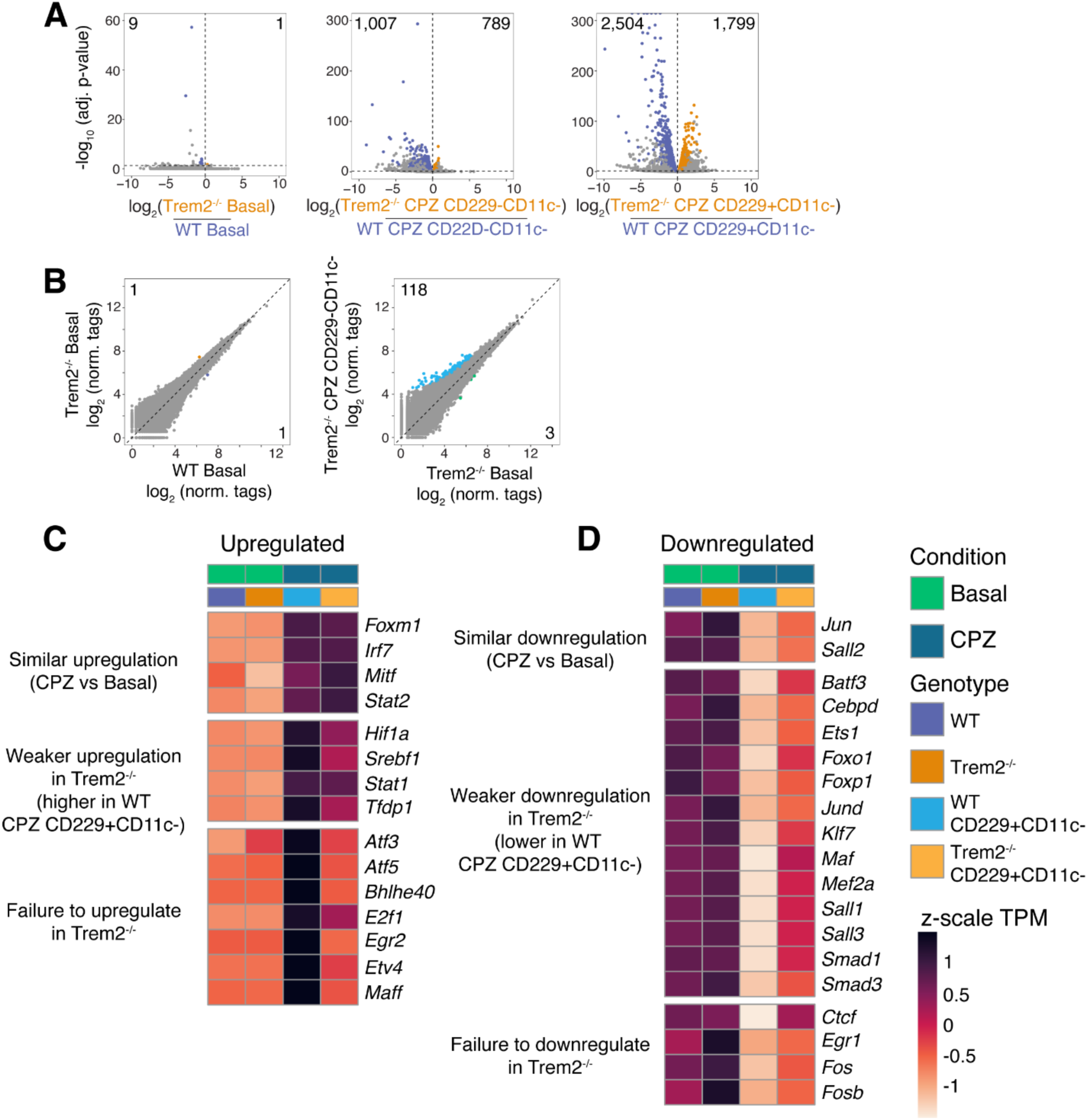
Differential gene expression analysis, comparing microglia from WT and Trem2^-/-^ fed regular or cuprizone diet for 4 weeks, related to Figure 3. **(A)** Volcano plots of RNA-seq data comparing microglia subsets isolated from WT and Trem2^-/-^ fed regular or CPZ diet for weeks. Differentially expressed genes are colored coded (DESeq2, FDR ≤ 0.05; *n* = 3-4). **(B)** Scatterplots of ATAC-seq data, centered on promoter-distal genomic sites, comparing profiles from homeostatic Trem2^-/-^ microglia subsets to WT counterpart (left) and Trem2^-/-^ CPZ CD229+CD11c- to Trem2^-/-^ basal microglia (right). Sites differentially accessible are colored coded (DESeq2, FDR ≤ 0.05; *n* = 2-3). **(C, D)** Heatmaps depicting z-scores of mRNA expression, comprising significantly modulated transcription factor genes in WT CPZ microglia as defined in Figure 2, comparing homeostatic and CPZ CD229+CD11c- from either WT or Trem2^-/-^ mice. Differentially expressed genes assessed with DESeq2 (FDR ≤ 0.05; *n* = 3-4).

**Supplementary Figure 4.**
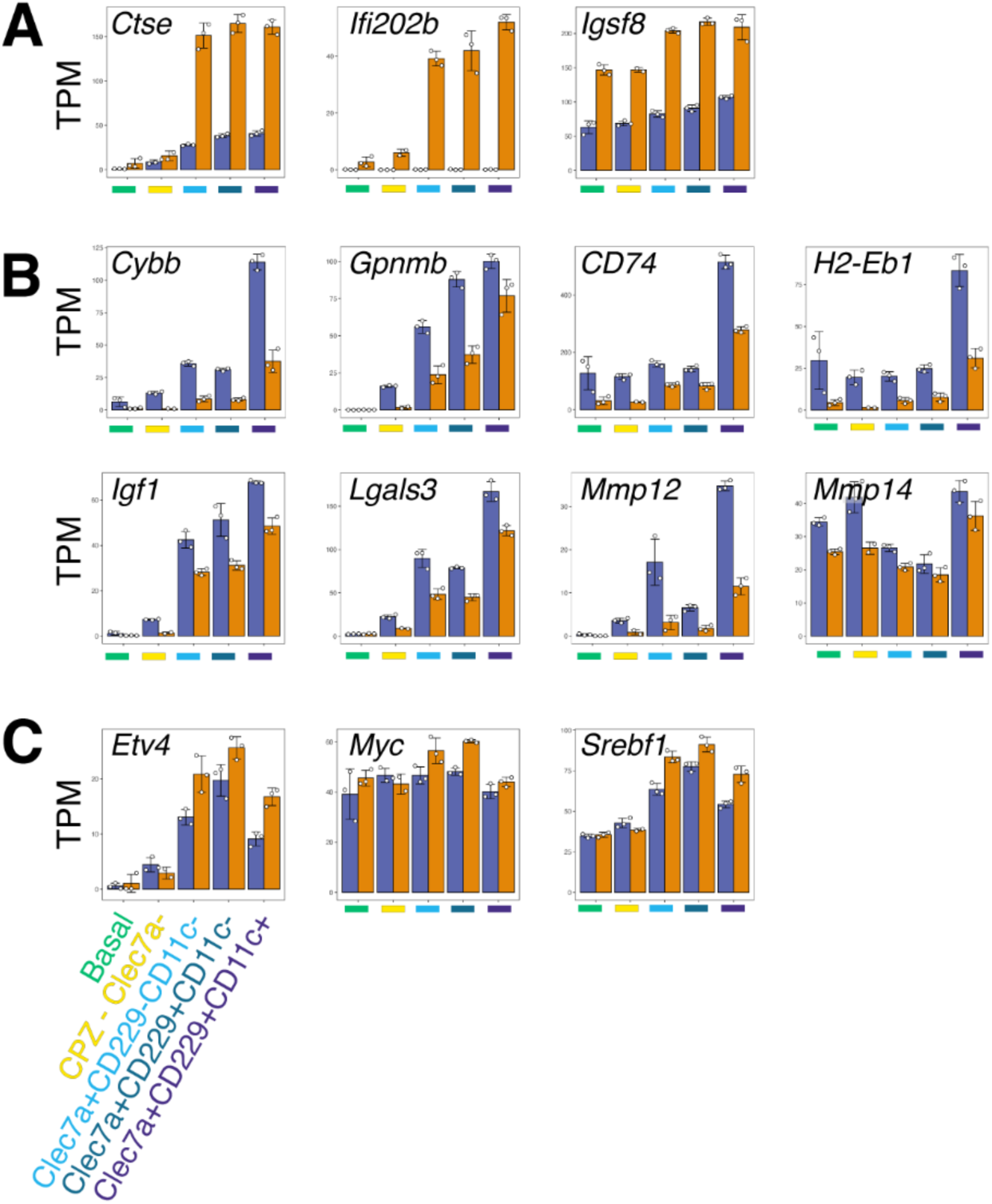
Bar graphs depicting mRNA expression of select genes preferentially expressed in distinct microglia subsets from Ctrl and MG-Mef2aΔ fed regular or cuprizone diet for 5 weeks, related to Figure 5. **(A)** mRNA expression levels (TPM) of select genes expressed at significantly higher levels in MG-Mef2aΔ CPZ subsets vs Ctrl counterparts. **(B)** mRNA expression levels (TPM) of select genes expressed at significantly higher levels in Ctrl CPZ subsets vs MG-Mef2aΔ counterparts. **(C)** mRNA expression levels (TPM) of select transcription factors expressed at significantly higher levels MG-Mef2aΔ CPZ subsets vs Ctrl counterparts.

**Table S1.** RNA-seq data (TPM) for genes differentially expressed in homeostatic and CPZ inflammatory microglia subset isolated from WT mice, related to figure 1.

**Table S2.** RNA-seq data (TPM) for transcription factors significantly upregulated or downregulated in CPZ inflammatory microglia compared to homeostatic microglia, related to figure 2.

**Table S3.** RNA-seq data (TPM) for genes differentially expressed in homeostatic and CPZ inflammatory microglia subset isolated from WT and Trem2^-/-^ mice, related to Figure 3.

**Table S4.** RNA-seq data (TPM) for genes differentially expressed in homeostatic microglia subset isolated from Ctrl and MG-Mef2aΔ mice, related to Figure 4.

**Table S5.** RNA-seq data (TPM) for genes differentially expressed in CPZ inflammatory microglia subset isolated from Ctrl and MG-Mef2aΔ mice, related to Figure 5.

**Table S6.** RNA-seq data (TPM) for genes differentially expressed in homeostatic and CPZ inflammatory microglia subset isolated from Ctrl and MG-Egr2Δ mice, related to Figures 6.

